# Liquid-liquid phase separation of intrinsically disordered FG-Nups is driven by highly-dynamic hydrophobic FG-motifs

**DOI:** 10.1101/2022.09.20.508740

**Authors:** Maurice Dekker, Erik van der Giessen, Patrick R. Onck

## Abstract

The intrinsically disordered FG-Nups in the NPC central channel can undergo liquid-liquid phase sepration (LLPS) into liquid condensates that display NPC-like permeability barrier properties. Here we present LLPS characteristics of each of the disordered FG-Nups of the yeast NPC. Using molecular dynamics simulations at amino acid resolution, FG-Nup condensates are studied and the main physicochemical driving forces for FG-Nup LLPS are identified. We show that FG-motifs that are predominantly present in the disordered domain of FG-Nups act as highly-dynamic hydrophobic stickers that are essential for the formation of stable liquid-like condensates. Next to that, we study LLPS of an FG-Nup mixture that resembles the NPC stoichiometry and observe that an NPC condensate is formed containing multiple types of FG-Nups. We find that the LLPS of this NPC condensate is also driven by FG–FG interactions, similar to the homotypic FG-Nup condensates. Based on the observed LLPS behavior, we categorize the different FG-Nups of the yeast NPC into two classes: The GLFG-Nups located in the central channel of the NPC phase separate into liquid-like condensates, forming a high-density cohesive barrier that can exclude inert particles. The FG-Nups at the entry and exit of the NPC channel, containing no GLFG-motifs, do not phase separate and possibly form a repulsive barrier by entropically excluding inert particles.

## Introduction

Nuclear pore complexes (NPCs) are the sole gateways for molecules to transit between the cytoplasm and the nucleus. NPCs are large protein complexes that have a highly selective permeability barrier, allowing small molecules to pass through passive diffusion, while the transport of larger molecules is significantly reduced, unless they are bound to nuclear transport receptors (NTRs). The central channel of the NPC is lined with intrinsically disordered nucleoporin proteins containing many phenylalanine/glycine (FG) motifs, i.e. FG-Nucleoporins (FG-Nups). These FG-motifs have been shown to play an essential role in NTR-driven transport [1, 2] and are also suggested to be important in maintaining the passive permeability barrier of the NPC [3–5]. Various descriptions exist of the phase state of the FG-Nups in the NPC central channel: The selective phase model [6, 7] proposes that the FG-Nups form a dense, sieve-like meshwork through specific FG–FG interactions, allowing small particles to pass, while blocking larger molecules. In the virtual gate model, or polymer brush model [8, 9], the NPC selectivity is ascribed to an entropic barrier formed by the FG-Nups, where small particles have a small entropic penalty and can easily cross the NPC, while large cargoes have a big entropic penalty and hence are less likely to traverse the NPC. The forest model [10] attributes the FG-Nup behavior to the bimodal distribution of charges along the FG-Nups sequences, allowing for molecules with an extended and a collapsed domain. This bimodal conformational character leads to a large variation of FG-Nup densities in side the NPC, resulting in different transport pathways for passive and NTR-driven transport. Finally, the Kap-centric model [11, 12] is based on the premise that NTRs are an integral part of the NPC and thus must play an important role in the pore’s barrier and transport function.

In recent years, liquid-liquid phase separation (LLPS) has been recognized to be a powerful mechanism of cells to organize their cytoplasm and nucleoplasm, and to catalyze molecular interactions through the formation of local high concentrations of interacting molecules [13, 14]. LLPS is omnipresent in cells and is deemed vital for a wide range of biological processes such as the formation of nucleoli [15, 16] and RNA-transcription [17–19]. LLPS of biomolecules has gained much attention, with a specific focus on RNA binding proteins (RBPs) and RNA-transcription [20–22]. Many RBPs contain low-complexity domains that are intrinsically disordered and can phase separate into liquid-like condensates, where the main physicochemical drivers have been identified to be cation–π interactions [22–25].

Experiments show that FG-Nups can also undergo phase separation into liquid condensates [26], hydrogels [7, 27–30] and even amyloid-like structures [31–33]. Most experiments specifically focus on the formation of FG-Nup hydrogels, as these show similar permeability properties as the NPC barrier [3, 7, 34]. Interestingly, recent work has shown that FG-Nup condensates that formed through LLPS are in a liquid-like state, which was found to be enough to reconstitute the selectivity of the NPC permeability barrier [26], i.e. small cargo molecules can enter FG-Nup condensates by passive diffusion, while big cargo can only join the FG-Nup phase with the help of NTRs. These in-vitro results, together with the important role of FG-Nup condensation in NPC biogenesis in-vivo [35], point to LLPS of FG-Nups as an essential organizing principle utilized by the cell in facilitating NPC assembly and selective transport.

Similar to the low-complexity domains in RBPs, FG-Nup sequences also exhibit a relatively low complexity with FG-motifs being interspaced by mostly polar amino acids. Due to the low abundance of charged residues in FG-Nups, electrostatic and cation–π interactions are not predominant, suggesting that LLPS of FG-Nups is driven by different molecular interactions than in the well-characterized RBP low-complexity domains.

Over the past years, several experiments have tested for phase separation of FG-Nups, showing phase transitions for Nup116 [28, 33], Nup100 [28, 33], Nup145N [33], Nup49 [26] and Nsp1 [7, 27, 34, 36]. Next to that, Patel et al. [37] have used bead-immobilized FG-Nups at high concentrations mixed with soluble fluorescently labeled FG-Nups to characterize the low-affinity protein interactions of all FG-Nups in yeast. Their experiments demonstrate the presence of homotypic attractive interactions for the FG domains of Nup116, Nup100, Nup145N, Nup49, Nup57 and Nup42 both in vivo and in vitro, and the absence of such low-affinity interactions for Nup159, Nsp1, Nup1, Nup2 and Nup60. However, a complete assay evaluating the LLPS characteristics for all yeast FG-Nups and their different domains is missing. Here, we investigate LLPS of FG-Nups by carrying out coarse-grained molecular dynamics (CGMD) simulations using a one-bead-per-amino-acid (1-BPA) MD model [38–40]. This model has been used previously to study nuclear transport through the yeast NPC [39, 41–46] and biomimetic nanopores [47–50], and to analyze the LLPS of dipeptide repeat proteins [51]. Here, we show that several intrinsically disordered FG-Nups of the yeast NPC can phase separate into liquid-like condensates and that this phase transition is mainly driven by the highly dynamic hydrophobic FG-motifs that line the FG-Nup sequences. Finally, we study LLPS of an FG-Nup mixture that resembles the NPC stoichiometry and observe the formation of a highly dynamic condensate containing multiple types of FG-Nups. From intermolecular contact statistics of the different types of FG-Nups, we find that the LLPS of this NPC condensate is also driven by FG–FG interaction, similar to the homotypic FG-Nup condensates. Based on our findings, we divide the intrinsically disordered FG-Nups of the yeast NPC into two groups: FG-Nups that show LLPS and FG-Nups that do not. Combined with the location at which these FG-Nups are anchored to the NPC scaffold, we relate the observed phase behavior to the phase state of the NPC’s central channel.

## Results

### Selection of FG domains

This study is confined to those FG-Nup domains in the yeast NPC that contain FG-motifs and that are lacking any ordered secondary structures (see Figure 1). To select these domains, we first determined the disordered domain for each FG-Nup using three different disorder predictors to minimize the effect of bias and over/under-prediction of disorder [52]. The disorder prediction scores show significant differences, but generally agree well on the location of the boundary between ordered and disordered domains. Kim et al. [53] also identified the disordered domains for each of the FG-Nups that constitute the yeast NPC. These domains are highlighted in Figure 1, where we considered both the intrinsically disordered and the flexible linker domains as part of the disordered domain. This choice is supported by the excellent agreement with the disorder prediction scores. Nup2 (not present in the data of Kim et al.) has an ordered domain at its C-terminal (AA 601–708) [54], in good agreement with the disorder prediction scores.

**Figure 1.**
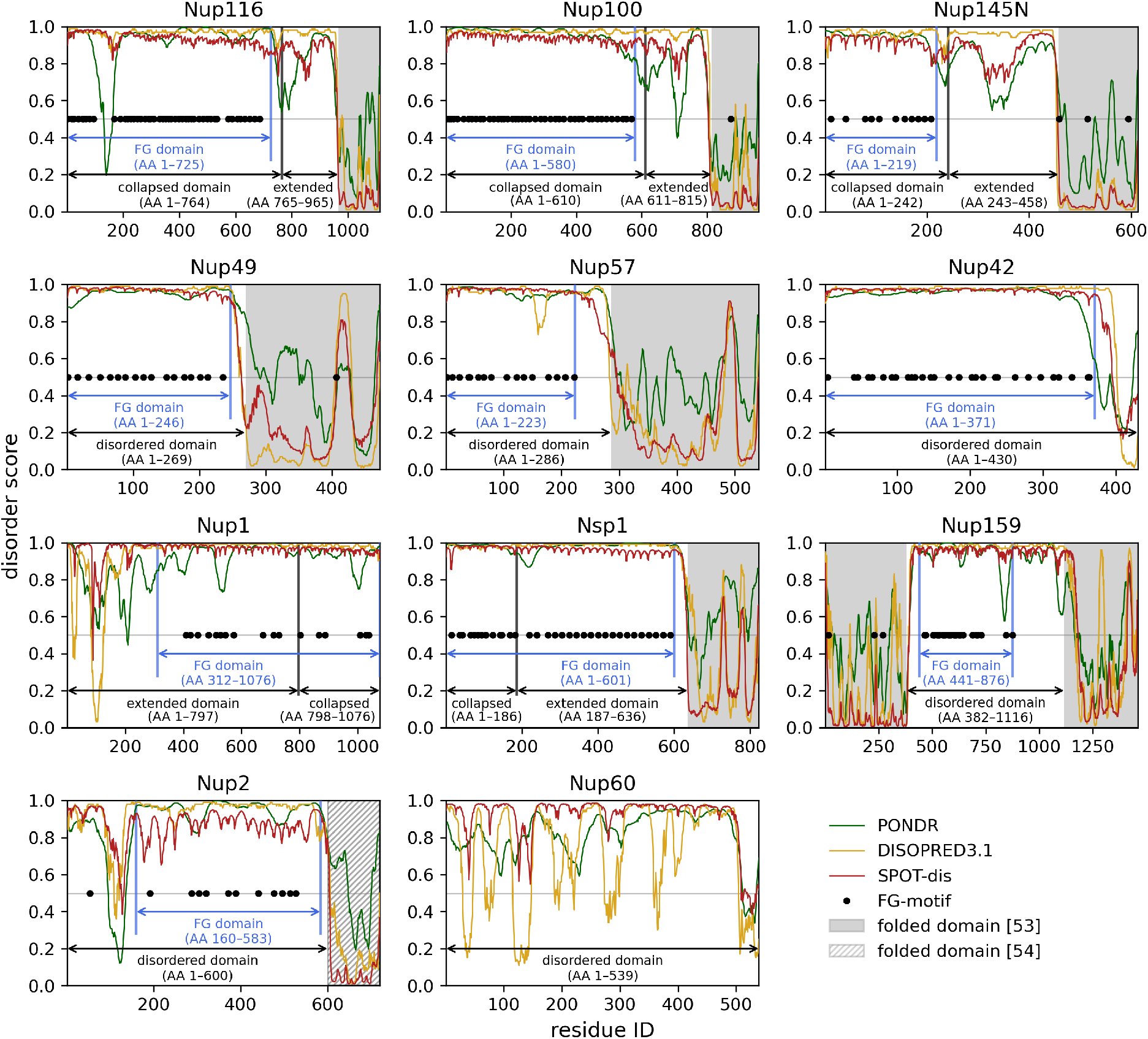
Selection of FG-Nup domains based on the intrinsic disorder and location of FG-motifs. Disorder prediction scores for the full-length FG-Nups are obtained from three different disorder predictors: PONDR [55], DISOPRED [56] and SPOT-Disorder [57]. Horizontal gray lines mark the boundary between disordered (scores > 0:5) and not disordered (scores < 0:5). Shaded areas indicate folded domains as previously identified by Kim et al. [53] and are in excellent agreement with the disorder prediction scores. Black dots mark the position of the FG-motifs in the sequence. The selected FG domains that are used in our simulations are marked by blue arrows, the disordered domains by black arrows. For the FG-Nups with a bimodal charge distribution (i.e. Nup116, Nup100, Nup145N, Nup1 and Nsp1 [10]), the collapsed and extended sections of the full disordered domain are indicated explicitly.

Having identified the full disordered domains of the FG-Nups, we selected the FG domains that contain all FG-motifs from the disordered domain (Figure 1, blue domains). Unless otherwise stated, these are the domains that are used in our simulations.

### LLPS of FG-Nups is driven by FG-motifs

To study the LLPS characteristics of FG-Nups, we evaluate the phase separation behavior of the FG domain of all FG-Nups (indicated in blue in Figure 1) using 1-BPA molecular dynamics simulations. To speed up the equilibration process, all simulations are initiated from a high-density protein condensate structure (see Figure 2, starting configuration). Within 2 µs of simulation time, all systems have reached a dynamic equilibrium state (Figure S2), which is either a stable liquid condensate surrounded by a low concentration of free monomers in the case of LLPS, or a homogeneous monomer solution when there is no LLPS. See the Methods section for more details.

**Figure 2.**
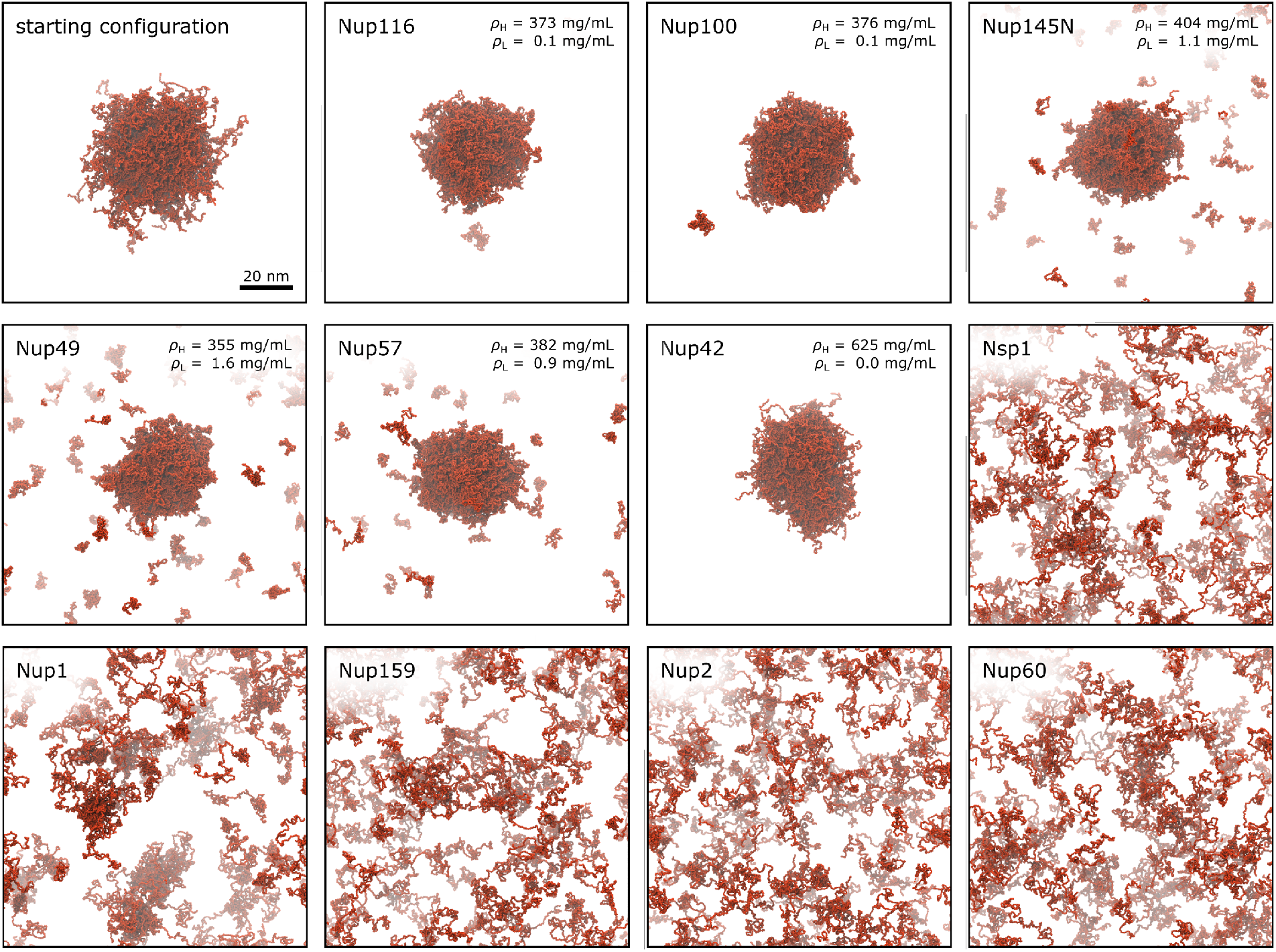
Overview of the LLPS behavior of each FG-Nup. An example of a starting configuration (Nup49) that is used for each simulation is shown next to simulation snapshots of each FG-Nup system at *t*^∗^ = 5 µs. LLPS is observed for the FG domains of Nup116, Nup100, Nup145N, Nup49, Nup57 and Nup42. For the systems where a stable condensate exists, the concentrations of the dense phase and dilute phase, *ρ*_H_ and *ρ*_L_, determined from radial density distributions (Figure S3) are shown in the top right corner of each snapshot.

Stable FG-Nup condensates are observed for Nup116, Nup100, Nup145N, Nup49, Nup57 and Nup42 (Figure 2). Each of these condensates is in dynamic equilibrium with the surrounding solvent, meaning that the rate at which molecules leave the condensate is equal to the rate at which molecules enter the condensate (see Figure S2 and Movie S1). Using radial density distributions, we have determined the concentrations of the dense phase and the dilute phase for each of the stable FG-Nup condensates (Figure S3). Apart from Nup42, the density in each of the FG-Nup condensates, *ρ*_H_, is comparable (350–400 mg/mL). Surprisingly, the dense concentration for condensates with practically no dilute phase (i.e. Nup100 and Nup116) is slightly lower than for some condensates where a dilute phase is present (i.e. Nup57 and Nup145N). This suggests that there is no relation between the concentrations of the dense phase, *ρ*_H_, and the concentration of the dilute phase, *ρ*_L_. On the other hand, that the concentration of the dilute phase is significantly higher for the shorter FG-Nups (i.e. Nup49, Nup57 and Nup145N), suggesting that the FG-Nup length also plays an important role in the phase behaviour, as shown before for dipeptide repeat proteins [51].

In order to asses which residues are driving the LLPS of FG-Nups, we analyzed the intermolecular contact statistics in each of the FG-Nup condensates. We first determined the time-averaged number of intermolecular contacts per protein replica as a function of residue number, resulting in ‘heat maps’ of high and low interacting regions along the sequences of each FG-Nup (Figures S4 and S5). Because the different FG-Nups drastically vary in size, the absolute contact numbers are not directly comparable between FG-Nup systems, but do highlight the most important interacting regions of each sequence. To be able to quantitatively compare the intermolecular contacts independently of the protein chain length, we summed the contacts over all contact pairs for each residue (Figure 3a). These one-dimensional contact summations display the number of inter-residue contacts each individual residue makes. Upon marking the location of the FG-motifs, we observe that many of the high contact peaks along the chains perfectly align with the location of FG-motifs. This shows that most of the intermolecular contacts in the FG-Nup condensates are with FG-motifs and that the high intensity spots in the full contact maps represent FG-FG interactions. These high contact probabilities for FG-motifs are observed in intramolecular contact maps of single-chain simulations as well, also in the case of FG-Nups that do not phase separate (Figure S8). This suggests that the presence of FG-motifs alone is not enough for FG-Nups to undergo LLPS.

**Figure 3.**
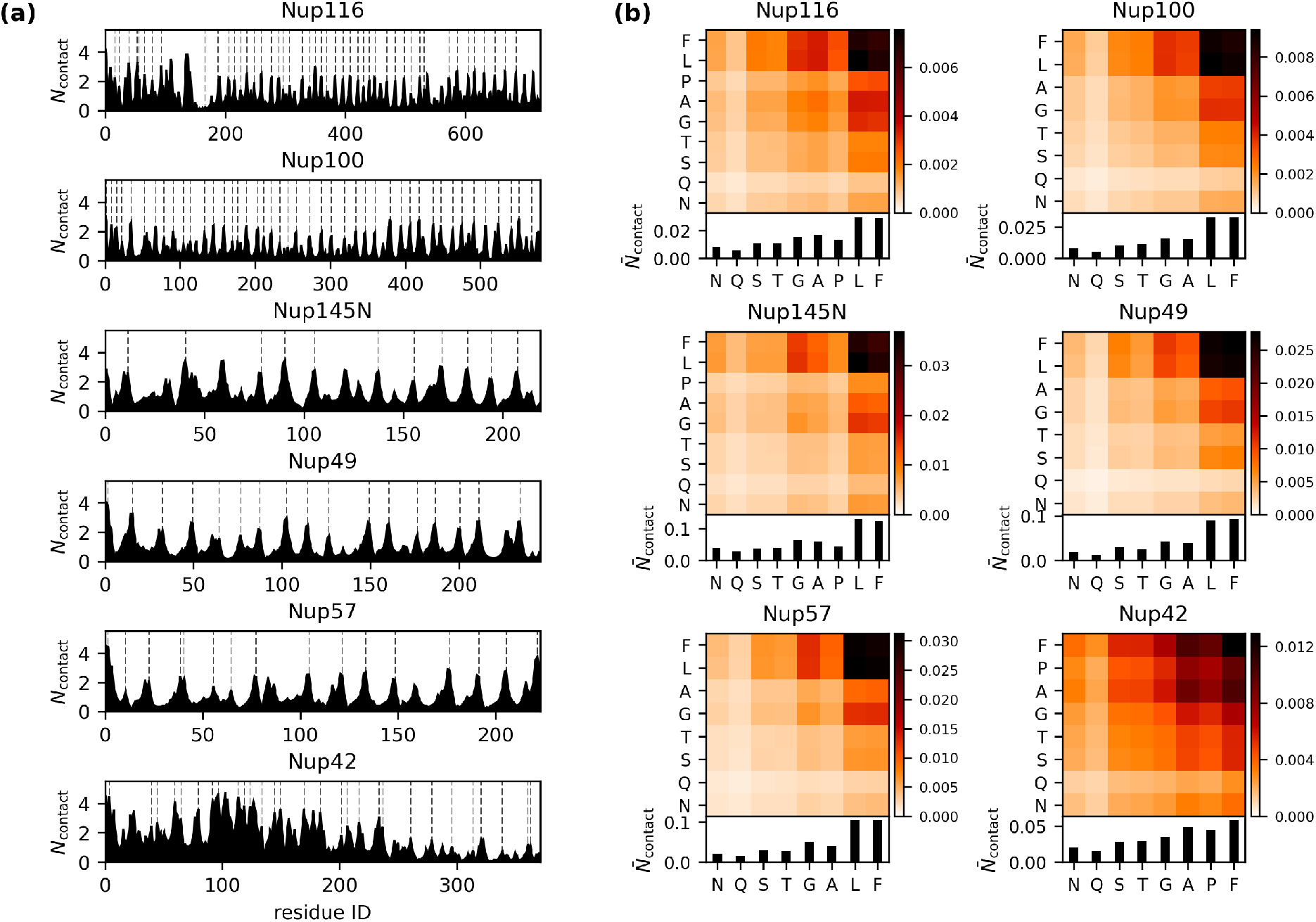
Characterizing intermolecular interactions in FG-Nup condensates. (**a**) Time-averaged number of intermolecular contacts per protein replica in the condensed phases of Nup116, Nup100, Nup145N, Nup49, Nup57 and Nup42 as a function of residue number. The dashed lines indicate the location of FG-motifs. Full contact maps showing the intermolecular contacts per residue pair are shown in Figures S4 and S5. (**b**) Time-averaged number of intermolecular contacts per protein replica in the condensed phases of Nup116, Nup100, Nup145N, Nup49, Nup57 and Nup42 as a function of residue type. The contact maps are normalized for residue abundance and only residues that contribute at least 4 % to the amino acid sequence are shown. The top panels show the (normalized) number of contacts between any pair of residue type; the bottom panels show the summed contact numbers per residue type. The non-normalized contact maps can be found in Figures S4 and S5.

To further investigate the interactions that are stabilizing FG-Nup condensates, we next analyzed the time-averaged number of intermolecular contacts per protein replica as a function of residue type (Figure 3b). The contact maps are normalized for amino acid abundance. We find that the contact maps for each of the GLFG-Nups are highly similar with most intermolecular interactions involving Phe (F) and Leu (L) residues. This highlights the importance of both residues for FG-Nup LLPS, which is also supported by recent mutation studies that showed that mutating either F or L of GLFG-motifs into Ala (A) results in a loss of LLPS [30].

Finally, we find that in each of the FG-Nup condensates around 50 % of the FG-motifs is involved in FG–FG interactions (Figure S10). Surprisingly, these FG–FG interactions are highly dynamic, with FG–FG contacts only existing for ∼ 1 ps in our simulations (Figure S11). After correcting for the coarse-grained dynamics speed-up (see SI Appendix), the lifetime of FG–FG contacts follows to be 0:03–0:1 ps in actual time. Nevertheless, the FG–FG contact lifetime is still significantly longer than those of other inter-residue contacts such as S–S or A–A.

### LLPS of bimodal FG-Nups

In the simulations above, we have used the FG domain of the FG-Nups (blue domains in Figure 1). However, several FG-Nups display a bimodal distribution of charged amino acids in their disordered domains, resulting in a collapsed and an extended domain [10]. These domains are indicated as well in Figure 1, following the definition for the collapsed/extended domains of Yamada et al. [10]. Because of the ascribed importance of the bimodal character of the FG-Nups to the selectivity of the NPC [10], we also investigated the role of bimodal charge distribution on the phase behavior of the different FG-Nups.

First of all, three of the central channel GLFG-Nups (Nup116, Nup100 and Nup145N) exhibit a clear bimodal charge distribution in their amino acid sequence [10]. Because the extended (stalk) domains of these FG-Nups are lacking FG-motifs, these domains were not included in the FG domains used above (Figures 2 and 3). However, considering their significant length (∼ 200 residues), it is reasonable to assume that these stalk domains alter the phase behavior of these FG-Nups. We have therefore repeated the LLPS simulations of Nup116, Nup100 and Nup145N, now including the extended stalk domains (from hereafter named Nup116+, Nup100+ and Nup145N+, respectively). When setting up these simulations in a similar manner as for the FG domains, we noticed that the repulsive extended domains break up the initial condensate structure, forming smaller micelle-like condensates with the non-cohesive extended domains residing on the surface of the condensates. With time these condensates started to coalesce, forming larger condensates. To speed up this process, we decided to build our initial condensate structure closer to the expected equilibrium state: a condensate structure with the cohesive collapsed domains in the center and the non-cohesive extended domains at the outside.

We have simulated the FG-Nup systems for at least 5 µs and observed that again the equilibrium state is reached within 2 µs (Figure S2). Phase separation is now only observed for Nup116+ and Nup100+, where both FG-Nup condensates show similar micelle-like conformations (see Figure 4a). In both simulations we observed a dynamic equilibrium state with a continuous exchange of molecules between the condensate and the dilute phase, showing that the initial conformation is not biasing the observed equilibrium state. We note that the dilute concentration for both Nup116+ and Nup100+ are slightly higher than for the respective FG domain condensates. On the other hand, the densities of the Nup116+ and Nup100+ condensates are slightly lower than the FG domain case (see Figure 4a and S3).

**Figure 4.**
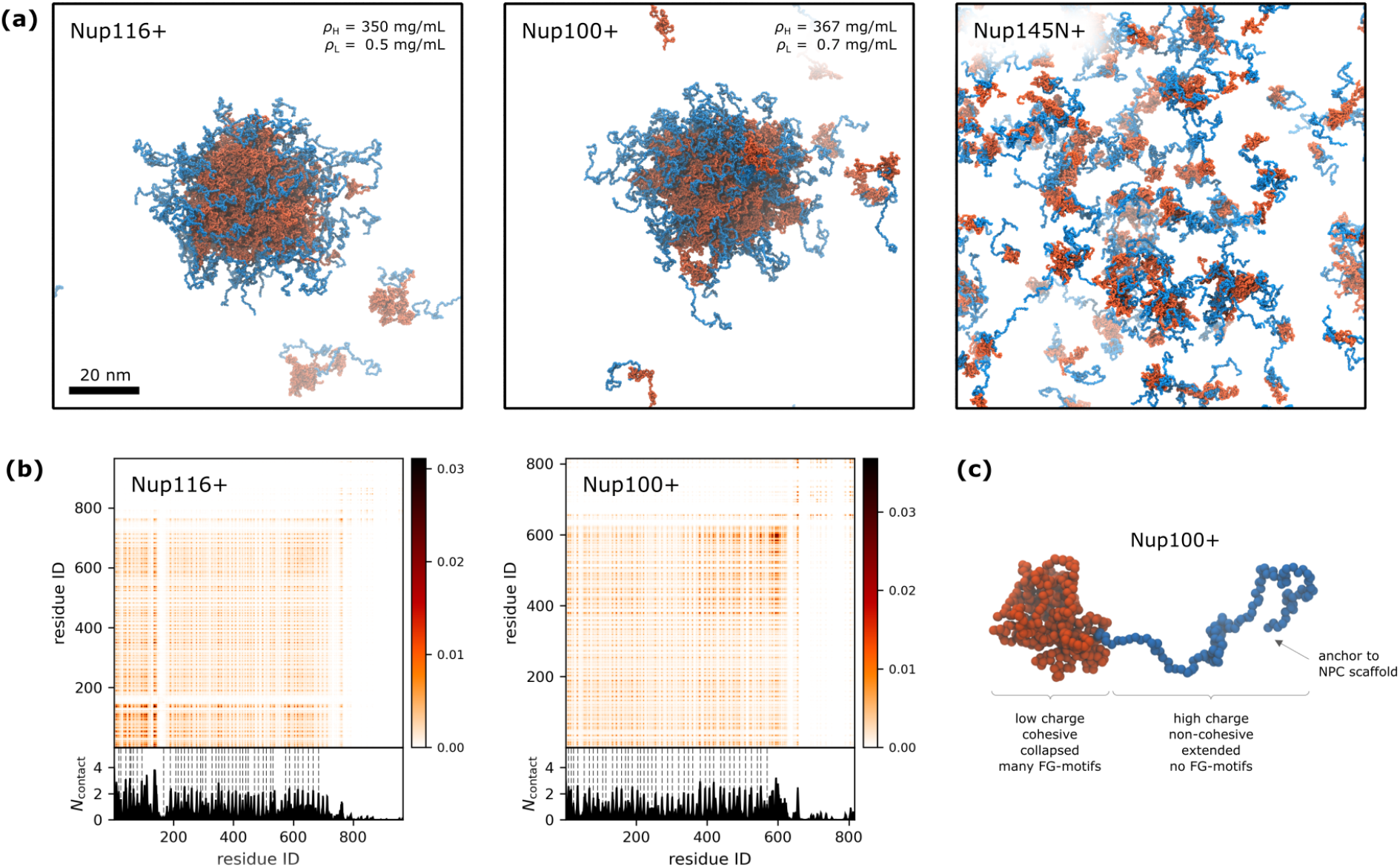
LLPS of the bimodal Nup116+, Nup100+ and Nup145N+. (**a**) Snapshots of the equilibrium states at *t*^∗^ = 5 µs. LLPS is observed for Nup116+ and Nup100+, where both condensates display a micelle-like structure: cohesive FG domains (red) form the core of the condensate with the extended stalk domains (blue) on the outside. No LLPS is observed for Nup145N. For Nup116+ and Nup100+ the concentrations of the dense phase and dilute phase, *ρ*_H_ and *ρ*_L_, determined from radial density distributions (Figure S3) are shown in the top right corner of the snapshots. (**b**) Time-averaged number of intermolecular contacts per protein replica in the condensed phases of Nup116+ and Nup100+ as a function of residue number. The dashed lines indicate the location of FG-motifs. The complete contact analysis of Nup116+ and Nup100+ can be found in Figure S6. (**c**) The bimodal character of Nup100+. The cohesive collapsed domain (red) contains many FG-motifs, the non-cohesive extended domain (blue) is devoid of any FG-motifs.

The intermolecular contact maps for the Nup116+ and Nup100+ condensates confirm that the extended domains have only little interactions with the collapsed domains (Figure 4b). Moreover, we note that the intermolecular contacts of the collapsed domain are very similar to those in the FG domain condensates (Figure S4).

Nup145+ did not display LLPS (Figure 4a), which might be caused by two factors: First of all, we note that the FG domain of Nup145N has a lower phase separation propensity then the FG domains of Nup116 and Nup100, suggested by the higher concentration of the dilute phase of Nup145N. Next to that, we also note that the cohesive domain of Nup145N is significantly shorter than the cohesive domain of Nup116 and Nup100, while the lengths of the stalk domains are comparable for all three FG-Nups. Clearly, the difference in length ratios of the collapsed and extended domains plays an important role in LLPS, which warrents further investigation.

### The collapsed domains of Nsp1 and Nup1 do phase separate

Besides Nup116, Nup100 and Nup145N discussed in the previous section, also Nsp1 and Nup1 features a bimodal charge distribution in their sequence (see Figure 1). Rather than a collapsed domain with FG-motifs and an extended domain devoid of FG-motifs, for Nsp1 and Nup1 the bimodality is within their FG domain, with both domains containing a similar density of FG-motifs. In the foregoing simulations, we observed no LLPS for the FG domains of Nsp1 and Nup1. Although Nup1 did not form a large condensate as was observed for the GLFG-Nups and Nup42, the LLPS simulation of Nup1 does show small clusters (5–10 molecules, see Figure 2) which seem to be driven by the low-charge collapsed domains. Also, it has been shown experimentally that the collapsed domain of Nsp1 can phase separate into hydrogels [7]. To clarify the role of these domains, we also conduct LLPS simulations for the collapsed domains of Nsp1 and Nup1.

We find that the collapsed domains of Nsp1 and Nup1 both form stable condensates (see Figure 5). Despite the fact that the collapsed domain of Nsp1 mostly contains regular FG-motifs, its LLPS behavior is very similar to that of the GLFG-Nups, with concentrations of the dense phase and dilute phase comparable to those of GLFG-Nups. Intermolecular contact analysis (Figure S7) highlights the considerable amount of FG-interactions present in the Nsp1 condensate, driving the LLPS of this FG-Nup segment.

**Figure 5.**
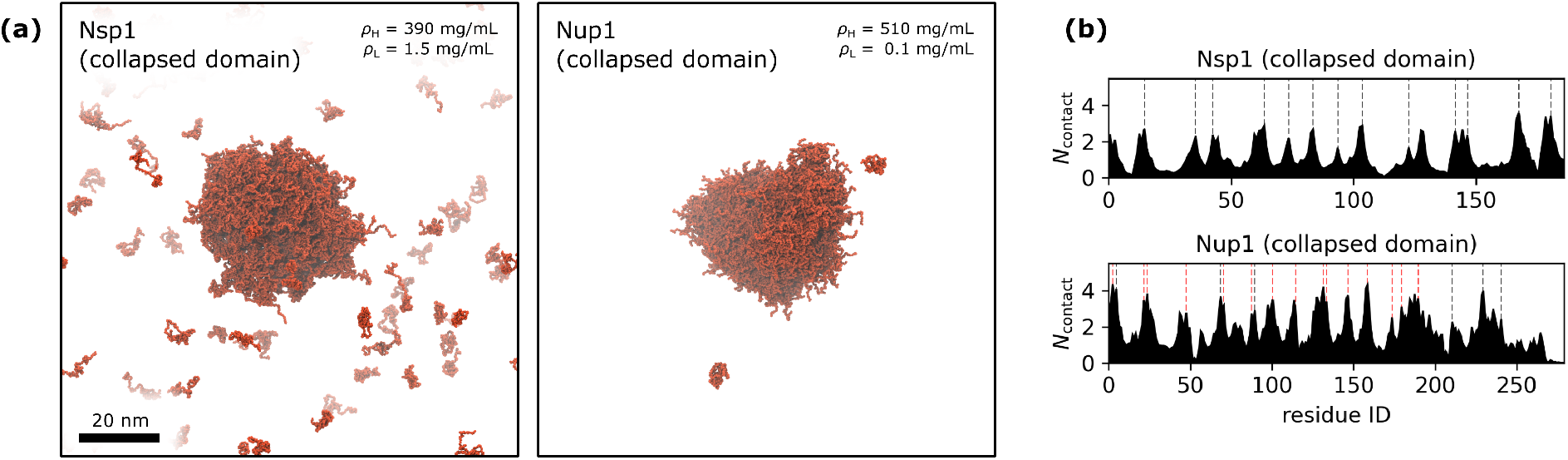
LLPS of the collapsed domains of Nsp1 and Nup1. (**a**) Snapshots of the equilibrium states at *t*^∗^ = 5 µs. The concentrations of the dense phase and dilute phase, *ρ*_H_ and *ρ*_L_, determined from radial density distributions (Figure S3) are shown in the top right corner of both snapshots. (**b**) Time-averaged number of intermolecular contacts per protein replica in the condensed phases of Nsp1 and Nup1 as a function of residue number. The black dashed lines indicate the location of FG-motifs. For Nup1 the location of F residues are indicated by red dashed lines. Full contact maps showing the intermolecular contacts per residue pair are shown in Figures S7.

The collapsed domain of Nup1 contains only six FG-motifs, yet phase separates into a stable FG-Nup condensate. Intermolecular contact analysis again confirms the presence of FG-interactions, but also reveals that other F residues (not part of an FG-motif) are involved in many of the intermolecular contacts (see Figure 5b, red dashed lines). This highlights that not only FG-motifs drive the phase separation of FG-Nups, but a significant amount of hydrophobic residues in general can, in line with the work of Najbauer et al. [30]. We note that the Nup1 condensate is not spherical as most of the other simulated FG-Nups condensates, but rather has a more ellipsoidal shape (see Figure 5). Nevertheless, there is a significant exchange of monomers between the condensate and the dilute phase. The radial density distribution of the Nup1c-condensate suggests a significantly higher density (see Figures 5 and S3). This high concentration of the dense phase might indicate the significant cohesiveness of the collapsed domain of Nup1, likely explaining the small clusters that we observed in the Nup1 FG domain simulation (Figure 2).

### Multiphase organization of FG-Nups in an NPC condensate

In the previous sections we have characterized the LLPS behavior of homotypic FG-Nup systems. Here, we will analyze LLPS for a mixed FG-Nup system to study how the presence of different types of FG-Nups (with different phase separation propensities) can modulate the collective LLPS behavior. The composition of the FG-Nup mixture is based on the stoichiometry of the yeast NPC [53], based on the full disordered domain of each FG-Nup (see Figure 1). By doing so, we test the latent phase state of the disordered domain of the NPC by releasing the FG-Nups from their anchor points on the NPC scaffold. As shown in Figure S12, the scaling exponents (see Supplementary Information) of the full disordered FG-Nup domains are not substantially different from the scaling exponents of the FG-Nups studied so far, justifying our choice to used the full disordered domains.

For the initial configuration of this NPC condensate simulation, we have build an FG-Nup condensate consisting of all FG-Nups with a scaling exponent below 0.5, i.e. the FG-Nups that are likely to phase separate. The other FG-Nups with scaling exponents larger than 0.5 are not expected to phase separate and are randomly placed in the dilute phase of the initial configuration (Figure S13). This approach increases the stability of the initial condensate, guaranteeing that during the simulation there is always a single condensate present. As a consequence of starting closer to the expected stable state, the dynamic equilibrium is reached significantly faster (see Methods for more details).

We have simulated the NPC condensate system for 10 µs, where a dynamic equilibrium is reached within 3 µs of simulation time (Figure S14). We observe a stable multiphase condensate accommodating the GLFG-Nups and Nup42 in different fractions (Figure 6a and Movie S3). The NPC condensate has an average diameter of approximately 35 nm, which is smaller than the inner diameter of the scaffold structure of the yeast NPC [53, 58, 59]. Similar to the homotypic LLPS simulations of Nup100+ and Nup116+, we again see that the non-cohesive extended domains of Nup100, Nup116 and Nup145N (colored in pink in Figure 6) mostly reside on the periphery of the condensate.

**Figure 6.**
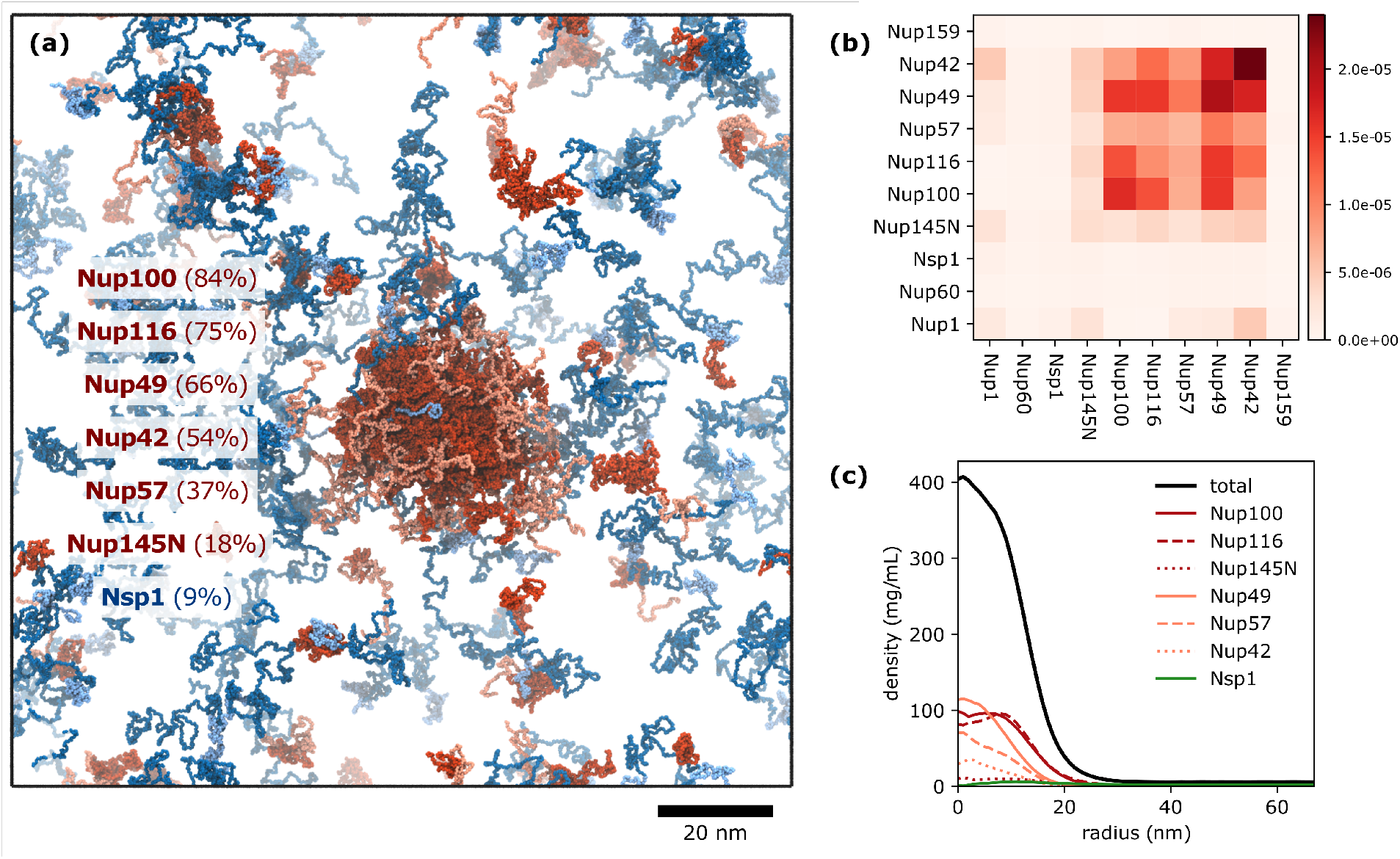
Analysis of the NPC condensate. (a) Snapshot of the equilibrium state of the NPC condensate at *t*^∗^ = 10 µs. FG-Nups with a scaling exponent *ν* < 0:5 are colored in red, with the extended domains of Nup116, Nup100 and Nup145N colored in pink. FG-Nups with a scaling exponent *ν* > 0:5 are colored in blue, with the collapsed domains of Nsp1 and Nup1 colored in light blue. The FG-Nups making up the NPC condensate are reported inside the snapshot, along with the fraction of each FG-Nup that is found in the dense phase, averaged over the last 5 µs of the simulation (Figure S14). (b) Time-averaged number of intermolecular contacts per protein replica for all FG-Nup combinations. The contact numbers are normalized for FG-Nup abundance. (c) Radial density distribution of the NPC condensate for the whole system and the most abundant FG-Nups in the dense phase.

The overall LLPS behavior of the different FG-Nups in the NPC condensate simulation is very similar to the homotypic LLPS simulations. However, there is a large variation in the phase separation propensity of the different FG-Nup types, highlighted by the different fractions of each FG-Nup present in the condensed phase. Most of the Nup100, Nup116 and Nup49 molecules are found in the condensed phase (70–90 %), while only half of the Nup57 and Nup42 molecules are in the NPC condensate (40–50 %). Surprisingly, where in the homotypic case the full disordered domain of Nup145N did not phase separate, here a notable fraction of Nup145N (20 %) is present in the condensed phase, showing that the phase separation propensity of FG-Nups can be modulated by the presence of other (phase separating) FG-Nups.

From the homotypic LLPS simulations we found that the FG domains of Nsp1 and Nup1 do not phase separate, while their collapsed domains do. In the NPC condensate we observe a significant amount of intermolecular contacts of the collapsed domains of Nsp1 and Nup1 with the FG-Nup condensate, although Nsp1 and Nup1 are not able to phase separate into the dense phase.

Intermolecular contact analysis on the NPC condensate (Figure 6b) reveals that there are interactions between each of the different FG-Nups in the NPC condensate. Most interactions are with Nup100, Nup49 and Nup42, which can be largely explained by the high abundance of these FG-Nups in the NPC condensate. The extensive amount of contacts with Nup42 is surprising considering only 50 % of the Nup42 monomers are in the condensed phase. Furthermore, the reduced amount of interactions with Nup57 and Nup145N can be explained by the lower occupancy of these FG-Nups in the NPC condensate. The full contact map, including intermolecular contacts per residue index, shows that the heterotypic interactions between FG-Nups are between the same domains as the homotypic interactions (Figure S15). In particular, we see that the extended domains of Nup100, Nup116 and Nup145N are almost completely devoid of any intermolecular interactions. On the other hand, a significant amount of contacts is observed for the collapsed domain of Nup1 (mostly with Nup42 and Nup145N present in the dilute phase). The similarity between the homotypic and heterotypic interactions supports our observations that LLPS of FG-Nups is driven by hydrophobic FG-motifs.

## Discussion

In this work, we used coarse-grained MD simulations at amino-acid resolution to study the LLPS of intrinsically disordered FG-Nups with the aim to identify the main physicochemical driving forces underlying the formation of FG-Nup condensates. We have tested the LLPS behavior over the full range of yeast FG-Nups and found that the FG domains of each of the GLFG-Nups and Nup42 phase separate into stable liquid-like condensates that are in dynamic equilibrium with their surroundings. No phase separation is observed for the FG domains of any of the FxFG-Nups and Nup159. Our results are in excellent agreement with in vitro experiments for both the phase separating and non-phase separating FG-Nups (see Table S4). Although the division in FG-motif type is striking, we note that the observed LLPS behavior is rather a result of the charge-to-hydrophobicity (C/H) ratio of the FG-Nup sequences. This is supported by the observed LLPS of Nup42, which is not of the GLFG-type, but does have a similar C/H value as the GLFG-Nups. Whereas charge–charge interactions are sometimes observed to drive the LLPS of polymers [60], the cationic and anionic residues in FG-Nups are homogeneously distributed along the sequences, resulting in a zero net charge along each of the FG domains.

Intermolecular contact analysis of the obtained FG-Nup condensates revealed the FG-motifs as the main drivers of LLPS of FG-Nups. This is in line with mutation studies that showed that mutating the phenylaline of the FG-motifs results in loss of phase separation [30, 37]. A closer look at the residues that are involved in the intermolecular interactions reveals the importance of hydrophobic residues for LLPS of FG-Nups. Besides the high contact numbers for F residues, we also find a similar amount of interactions with L residues (mostly part of GLFG-motifs), suggesting an equal importance of both F and L residues in the GLFG-motifs. This is supported by recent observations that mutating either the F or L of GLFG-motifs results in absence of LLPS of FG-Nups [30]. We note, however, that experiments have shown that interactions between aromatic residues (such as F–F interactions) are significantly stronger than normal hydrophobic interactions (such as L–L or L–F interactions) [61]. However, aromatic residues in our model merely interact through non-specific hydrophobic interactions and our model does not include specific π–π stacking or T-stacking interactions that might be present between the FG-motifs [30].

Besides the FG domains of each of the FG-Nups, we have also studied several other FG-Nup domain selections, specifically in the case of FG-Nups that have a bimodal charge distribution along their sequences. First of all, we have looked at the full disordered domains of Nup116, Nup100 and Nup145N. Apart from a cohesive FG domain, these FG-Nups also contain a highly charged extended domain that is devoid of FG-motifs. Upon including these extended domains into the FG-Nup sequences Nup100+ and Nup116+ still showed LLPS. The non-cohesive extended domains largely reside on the periphery of the condensates, forming micelle-like structures, and do not significantly participate in intermolecular interactions. We note that these extended domains connect the FG domains to the NPC scaffold [53, 54] and therefore hypothesize that a similar effect might take place in the central transporter channel of the NPC [39]. These non-cohesive domains then form low protein density areas between the NPC scaffold and central channel FG-meshwork, in correspondence with the forest model [10]. Additionally, we have also simulated LLPS of the low-charge cohesive domains of Nsp1 and Nup1. The collapsed domains of both these FG-Nups displayed LLPS, whereas the full-length FG domains did not. Our results are in line with the experiments of Frey et al. [7] that show that the collapsed domain of Nsp1 can phase separate (forming FG-hydrogels).

Finally, we studied the intermolecular interactions in a multiphase FG-Nup condensate with a stoichiometry similar to the yeast NPC. Here, we observed a stable GLFG-condensate, composed out of all the GLFG-Nups and Nup42. These are the same FG-Nups for which homotypic LLPS was observed, and we find that the same interactions are responsible for the phase separation of the NPC condensate. However, the dilute concentration of the NPC condensate is surprisingly high, with much higher fractions of free GLFG-Nups compared to the homotypic GLFG-Nup condensates.

We note that most of the FG-Nups for which we observed LLPS are located at the center of the yeast NPC (Nup116, Nup100, Nup145N, Nup49 and Nup57), while FG-Nups that do not phase separate are located at the cytoplasmic and nucleoplasmic sides of the NPC (Nup159, Nup1, Nup2 and Nup60). This suggests that the LLPS propensity of FG-Nups is related to the location and function in the NPC. Based on the above results, we hypothesize that the central channel FG-Nups form a liquid-like condensed phase, formed by highly-dynamic FG–FG interactions. The peripheral FG-Nups do not phase separate and form an entropic brush, blocking large inert molecules from entering the NPC, yet ignoring small particles that passively diffuse through the NPC.

## Methods

### Coarse-grained model for disordered proteins

All coarse-grained MD simulations are done with our 1-BPA model for intrinsically disordered proteins. This implicit-solvent model at amino-acid resolution samples from realistic backbone conformations and accounts for hydrophobic, electrostatic and cation–π interactions [38–40, 51]. Several hydrophobic interactions are fine-tuned to better match the experimental Stokes radii of yeast FG-Nup segments. All simulations are performed at 300 K and physiological salt concentration of 150 mM using the GRO-MACS [62] molecular dynamics software (version 2019.4), where the stochastic dynamics integrator operates with a time step of 0:02 ps and inverse friction coefficient *γ* ^−1^ = 50 ps. More details on the model and the improvements that have been made are provided in the Supporting Methods.

### FG-Nup LLPS simulations

To set up the system, we created a condensate structure by performing a 500 ns NVT equilibration in a cubic periodic box of at a density of 2 beads per nm^3^ (300–400 mg/mL), which is comparable to the densities of FG-Nup condensates (200–300 mg/mL [28]). Then, the periodic box dimensions are increased by 350 % with the protein condensate at the center (see Figure 2 for an example of the initial configuration). This results in an overall concentration of approximately 8 mg/mL. This initial configuration is preferred over a homogeneous monomer solution, as it significantly speeds up the equilibration process, while resulting in the same dynamic equilibrium state (i.e. a dense phase with a single FG-Nup condensate surrounded by a dilute phase with a low concentration of free monomers). For each FG-Nup system a system size was chosen such that the total number of amino acids is close to 70 000 beads, which is sufficiently large so that finite size effects in the simulations are small. The FG-Nup systems are simulated for 5 µs, where dynamic equilibrium is reached within the first 2 µs (Figure S2).

Cluster sizes (Figure S2) are determined using the *gmx clustsize* utility of GROMACS, where two monomers are considered to be in the same cluster if at least two residues of those monomers are within 0:7 nm [51].

### NPC condensate LLPS simulation

The LLPS simulation for the NPC condensate could be set up in a similar way as the homotypic FG-Nup LLPS simulations (i.e. starting from a condensate structure containing all FG-Nups). However, due to strong repulsive interactions from several non-cohesive FG-Nup domains (e.g. Nup159, Nup60, etc.), the initial condensate structure quickly falls apart and cohesive FG-Nups phase separate again into smaller condensate structures. Given enough time these smaller condensates will eventually merge into a single condensate, but given the large system size and the slow diffusion speed of the condensates, this might take tens of microseconds.

To prevent too strong repulsive interactions in the initial condensate structure, we have created an initial condensate structure containing only cohesive FG-Nups with scaling exponents below 0.5 (i.e. FG-Nups that are likely to phase separate: Nup116, Nup100, Nup145N, Nup49, Nup57 and Nup42; cf. Figure S12). The high-density condensate simulation is equilibrated for 500 ns (NVT), after which the box dimensions are increased and the non-cohesive FG-Nups with high scaling exponents are randomly distributed around the FG-Nup condensate, where the protein backbone conformations are sampled from single-chain simulations. The final box dimensions are chosen such that the concentration is the same as in the homotypic LLPS simulations. The resulting structure is a condensate containing all cohesive FG-Nups, surrounded by a homogeneous solution of non-cohesive FG-Nups (Figure S13). This initial configuration is then equilibrated for 10 µs, where the last 5 µs are used for analysis of the NPC condensate.

**Table 1.** Composition of the NPC condensate simulation, where the number of each FG-Nup is the same as in the yeast NPC [53]. For each FG-Nup the full disordered domain is used (see Figure 1). The FG-Nups are placed either in the dense phase or dilute phase of the initial configuration based on their cohesiveness (cohesive: *ν* < 0:5, non-cohesive: *ν* > 0:5; see Figure S12).

| FG-Nup | Domain | Frequency | Scaling exp. | Cohesive |
| --- | --- | --- | --- | --- |
| Nup116 | AA 1–965 | 16 | 0.49 | yes |
| Nup100 | AA 1–815 | 16 | 0.48 | yes |
| Nup145N | AA 1–458 | 16 | 0.49 | yes |
| Nup49 | AA 1–269 | 32 | 0.44 | yes |
| Nup57 | AA 1–286 | 32 | 0.46 | yes |
| Nup42 | AA 1–430 | 8 | 0.42 | yes |
| Nup1 | AA 1–1076 | 8 | 0.53 | no |
| Nsp1 | AA 1–636 | 48 | 0.57 | no |
| Nup159 | AA 382–1116 | 16 | 0.59 | no |
| Nup60 | AA 1–539 | 16 | 0.56 | no |

### Radial density profiles

Radial density profiles are obtained by determining the average density in discrete concentric shells of thickness 1 nm measured from the center of the condensate. The radial density profiles (Figures 6c and S3) are time-averaged over the last 3 µs of the simulation, where a density profile is determined every nanosecond. The concentration of the dense phase is obtained from the radial density profiles by averaging the density until the first point close to the high-density region where | *dρ* / *dr* | ≤ 2 − 5 mg/mL/nm [51] (see dashed lines in Figure S3). Similarly, the concentration of the dilute phase is obtained by averaging the density starting from the first point close the low-density region where | *dρ* / *dr* | ≤ 2 − 5 mg/mL/nm.

### Inter-residue contact analysis

Intramolecular contact analysis is performed on single-chain simulations. For each trajectory frame, an upper triangular contact matrix is calculated that describes all inter-residue contacts within the protein chain, where the three nearest neighboring residues along the chain are excluded. Here, any residue pair is considered to form a contact when the distance separating the beads is less than the cut-off value 0:7 nm. The average contact map is then calculated by normalizing for the length of the trajectory.

Intermolecular contact analysis is performed on LLPS simulations. For each trajectory frame, we first determined which molecules are part of the condensate using the *gmx clustsize* utility of GROMACS. Here, a protein chain is considered to be part of the condensate if at least one of the residues is within the cut-off distance of 0:7 nm of any bead of the condensate. Then, for all chains in the condensate, a contact matrix is calculated that describes all unique contacts between all residues in the condensate for that frame. This results in *N*_Nup_(*N*_Nup_ − 1)/2 contact matrices (where *N*_Nup_ is the number of FG-Nup proteins in the condensate) for each residue pair in the two FG-Nup molecules. The average number of intermolecular contacts by residue index are then obtained by summing all the individual contact matrices and normalizing for the number of chains in the condensate. Finally, these contact maps are averaged over the trajectory.

