## Supplementary Information for "Liquid-liquid phase separation of intrinsically disordered FG-Nups is driven by highly-dynamic hydrophobic FG-motifs"

### 1 Coarse-grained 1-BPA force field

The one-bead-per-amino-acid (1-BPA) model accounts for residue-specific hydrophobicity, charge and backbone stiffness. Proteins in the 1-BPA force field display realistic backbone conformations that are sampled from atomistic Ramachadran data [1]. The non-bonded interaction potential of the 1-BPA force field is a combination of hydrophobic, electrostatic and cation–$\pi$ terms. The attractive hydrophobic and repulsive hydrophilic interactions between residues are described by a shifted 8–6 Lennard-Jones potential:

$$\phi_{\text{hp}}(r) = \begin{cases} \varepsilon_{\text{rep}} \left(\frac{\sigma}{r}\right)^8 - \varepsilon_{ij} \left[\frac{4}{3}\left(\frac{\sigma}{r}\right)^6 - \frac{1}{3}\right] & \text{for} \quad r \leq \sigma, \\ (\varepsilon_{\text{rep}} - \varepsilon_{ij}) \left(\frac{\sigma}{r}\right)^8 & \text{for} \quad \sigma \leq r, \end{cases}$$

where $\sigma = 0.60\,\text{nm}$. The interaction strength for each pair of amino acids $(i, j)$ is given by the combination rule

$$\varepsilon_{ij} = \varepsilon_{\text{hp}} \sqrt{(\varepsilon_i \varepsilon_j)^{\alpha}},$$

where $\varepsilon_i$ is the relative hydrophobicity scale of residue $i$ (Table S1) and the exponent $\alpha = 0.27$ is chosen such that the model accurately reproduces experimental hydrodynamic radii of FG-Nup segments [2]. The variables $\varepsilon_{\text{hp}}$ and $\varepsilon_{\text{rep}}$ are chosen to be 13 and $10\,\text{kJ} \cdot \text{mol}^{-1}$, respectively, such that the minimum interaction energy of the most hydrophobic amino acids is $-5.22\,\text{kJ} \cdot \text{mol}^{-1}$.

Electrostatic interactions between charged amino acids are modelled with the modified Coulomb potential:

$$\phi_{\text{el}} = \frac{q_i q_j}{4\pi\varepsilon_0 \varepsilon_r(r) r} \exp(-\kappa r),$$

where $q_i$ is the charge of residue $i$ and $\varepsilon_0$ is the vacuum permittivity. The Debye screening coefficient is chosen to be $\kappa = 1.27\,\text{nm}^{-1}$, corresponding to a salt concentration of 150 mM. The dielectric screening of the solvent is given by the sigmoidal function

$$\varepsilon_r(r) = S_{\text{s}} \left[1 - \frac{r^2}{z^2} \frac{e^{r/z}}{\left(e^{r/z} - 1\right)^2}\right],$$

where $S_{\text{s}} = 80$, and $z = 0.25\,\text{nm}$.

In recent work, cation–$\pi$ interactions have been introduced to the 1-BPA force field [3]. Rather than interacting through the hydrophobic potential $\phi_{\text{hp}}$, cationic residues (e.g. R, K) interact with aromatic residues (e.g. F, Y, W) through an 8–6 Lennard Jones potential:

$$\phi_{\text{cp},ij} = \varepsilon_{\text{cp},ij} \left[3\left(\frac{r_{\text{m}}}{r}\right)^8 - 4\left(\frac{r_{\text{m}}}{r}\right)^6\right],$$

where $\varepsilon_{\text{cp},ij}$ are pair-specific interactions energies (provided in Table **??**) and $r_{\text{m}} = 0.45\,\text{nm}$ is the equilibrium interaction distance.

#### 1.1 Update of the 1-BPA force field

In general, the 1-BPA force field shows very accurate protein conformations, especially for important FG-Nups such as Nup100 and Nup116. However, we also noticed that several FG-Nup segments display significant errors in their predicted Stokes radii, especially Nup49 and the collapsed domain of Nsp1. Considering the importance of gyration radius on the observed LLPS behavior, we have tried to improve the 1-BPA force field even further by updating the hydrophobicity values for several highly abundant amino acids: glycine (G), glutamine (Q) and aspargine (N). Here, our goal was to maintain the minimal error in Nup100 and Nup116, while rebalancing the hydrophobicities such that the errors in Nsp1 and Nup49 are reduced. As a consequence, also the errors for Nup145N, Nup159 and Nup1 are significantly improved (see Figure S1 and Table S3). The updated hydrophobicity values for G, Q and N that are used in this paper are:

| | | | |
|---|---|---|---|
| G: | 0.41 | $\rightarrow$ | 0.48 |
| Q: | 0.64 | $\rightarrow$ | 0.33 |
| N: | 0.33 | $\rightarrow$ | 0.41 |

#### 2 Coarse-grained dynamics speed-up of 1-BPA simulations

The dynamics observed in CG simulations are generally much faster than those in all-atom simulations. One of the main reasons is the use of implicit solvents with high inverse friction coefficients $\gamma^{-1}$, resulting in the simulated molecules experiencing very low friction and stochastic forces [4]. Combined with the smoother energy landscape arising from the larger particle sizes (i.e. reduction of degrees of freedom), the dynamics speed-up in CG simulations can be tremendous.

A general approach for determining the CG dynamics speed-up factor is by comparing long time-scale properties, such as diffusion coefficients, between CG systems and all-atom systems [5–7]. In principle, the diffusion rate of a particle in a solvent is described by the Stokes–Einstein relation:

$$D_\mathrm{s} = \frac{k_\mathrm{B} T}{6\pi\eta R_\mathrm{s}}, \tag{1}$$

where $\eta$ is the viscosity of the solvent and $R_\mathrm{s}$ is the hydrodynamic (or Stokes) radius of the particle. However, as in many CG models, the 1-BPA force field assumes overdamped Langevin dynamics (i.e. $\gamma^{-1} = 50\,\mathrm{ps} \geq \Delta t = 0.02\,\mathrm{ps}$), meaning that the diffusion coefficient of a molecule only depends on its molecular mass, and instead is completely described by the Einstein relation:

$$D_\mathrm{s} = \frac{k_\mathrm{B} T}{m\gamma}, \tag{2}$$

where $T = 300\,\mathrm{K}$ and the inverse friction coefficient $\gamma^{-1} = 50\,\mathrm{ps}$.

To determine the dynamics speed-up of our simulations, we have calculated the simulation diffusion coefficients $D_\mathrm{sim}$ for a range of disordered and folded proteins using equation (2) and compared this to the experimentally determined diffusion coefficients $D_\mathrm{exp}$. The CG speed-up, defined as $D_\mathrm{sim}/D_\mathrm{exp}$, are shown in Figure S9 for a range of molecular masses. As a consequence of the overdamped Langevin dynamics, the diffusion rates in our simulations are independent of the shape of the molecules and rather are completely described by the molecular mass. The molecular weights of the FG-Nups used in this paper range from 21 kDa (Nup57) to 100 kDa (Nup116), so we suggest that the dynamics speed-up in our simulations is in the range of 30–90 times. Because of the large range of observed speed-up factors, we decided to report all times in this paper as simulation times, unless explicitly stated (denoted as effective time).

#### 3 Relating single-chain characteristics to large-scale LLPS behavior

It was suggested that protein LLPS behavior can be obtained from single-chain simulations [8, 9], which would be a much more effective method to test LLPS behavior for large sets of proteins. We use the calculated scaling exponents as a probe for phase separation, where the exponent of 0.5 corresponds to an ideal chain (the so-called Θ state) in which the effective interactions between monomers are on average neither repulsive nor attractive. For chains where the effective interactions are attractive, i.e. $\nu < 0.5$, one would expect a chain to phase separate. To verify that protein scaling exponents are also a reliable measure for characterizing LLPS of FG-Nups, we calculate scaling exponents for all FG-Nups used in our bulk simulations (see Methods section). The calculated scaling exponents for each of the FG domains used (black points in Figure S12) are in excellent agreement with the simulated phase separation behavior. All FG-Nup segments for which we found LLPS have exponents well below 0.5 and all FG-Nup segments that did not phase separate have exponents well above 0.5. The reduction in phase separation propensity when including the extended domains of Nup116, Nup100 and Nup145N is also clear in the calculated scaling exponents (red points in Figure S12). We note that the only outlier is the full disordered domain of Nup145N, that did not phase separate in our simulations but was expected to based on the calculated exponent. We suggest that the the protein scaling exponents overall are excellent indicators for LLPS of FG-Nup segments, however, care should be taken for exponents close to the boundary of $\nu = 0.5$. The calculated exponents suggest that our results are not affected by the choice of FG domains, as for each FG-Nup (apart from the bimodal Nup116, Nup100 and Nup145) the scaling exponent of the FG domain is very close to the exponent of the full disordered domain (see Figure S12).

##### 3.1 Determination of protein scaling exponents

All single-chain simulations are initiated from an extended conformation. A brief equilibration (5 ns) is followed by a 10 µs production run from which a sample is taken every 0.2 ns, resulting in 50 000 protein conformations that are used to compute the average radius of gyration, $R_\mathrm{g}$. The protein scaling exponent, $\nu$, is then calculated from the average radius of gyration using equation [8, 9]

$$R_\mathrm{g} = \sqrt{\frac{\gamma(\gamma+1)}{2(\gamma+2\nu)(\gamma+2\nu+1)}}\, bN^{\nu}, \tag{3}$$

where $\gamma \approx 1.1615$, $b = 0.55\,\mathrm{nm}$ is the Kuhn length for disordered proteins and $N$ is the number of beads per chain.

### 4 Supplementary Figures

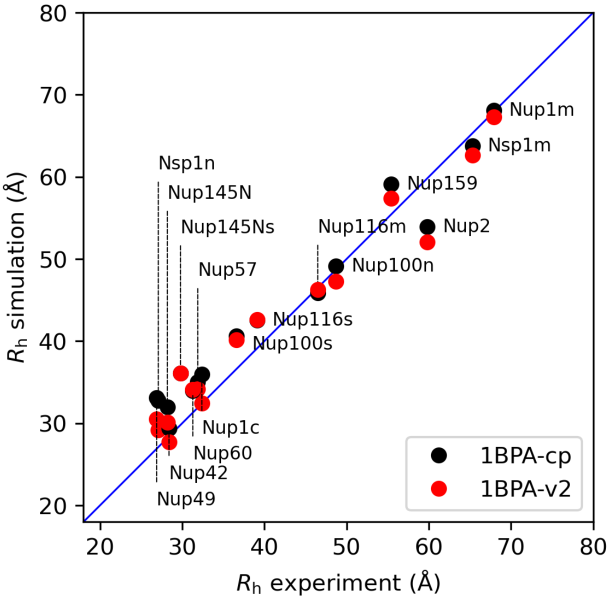

**Figure S1.** A direct comparison of the two force fields in predicting the hydrodynamic radius of FG-Nup segments. The total average and the largest errors are found to be 9.5 % and 23.1 % in the old 1-BPA-cp force field, and 6.5 % and 21.1 % in the updated 1-BPA-v2 force field.

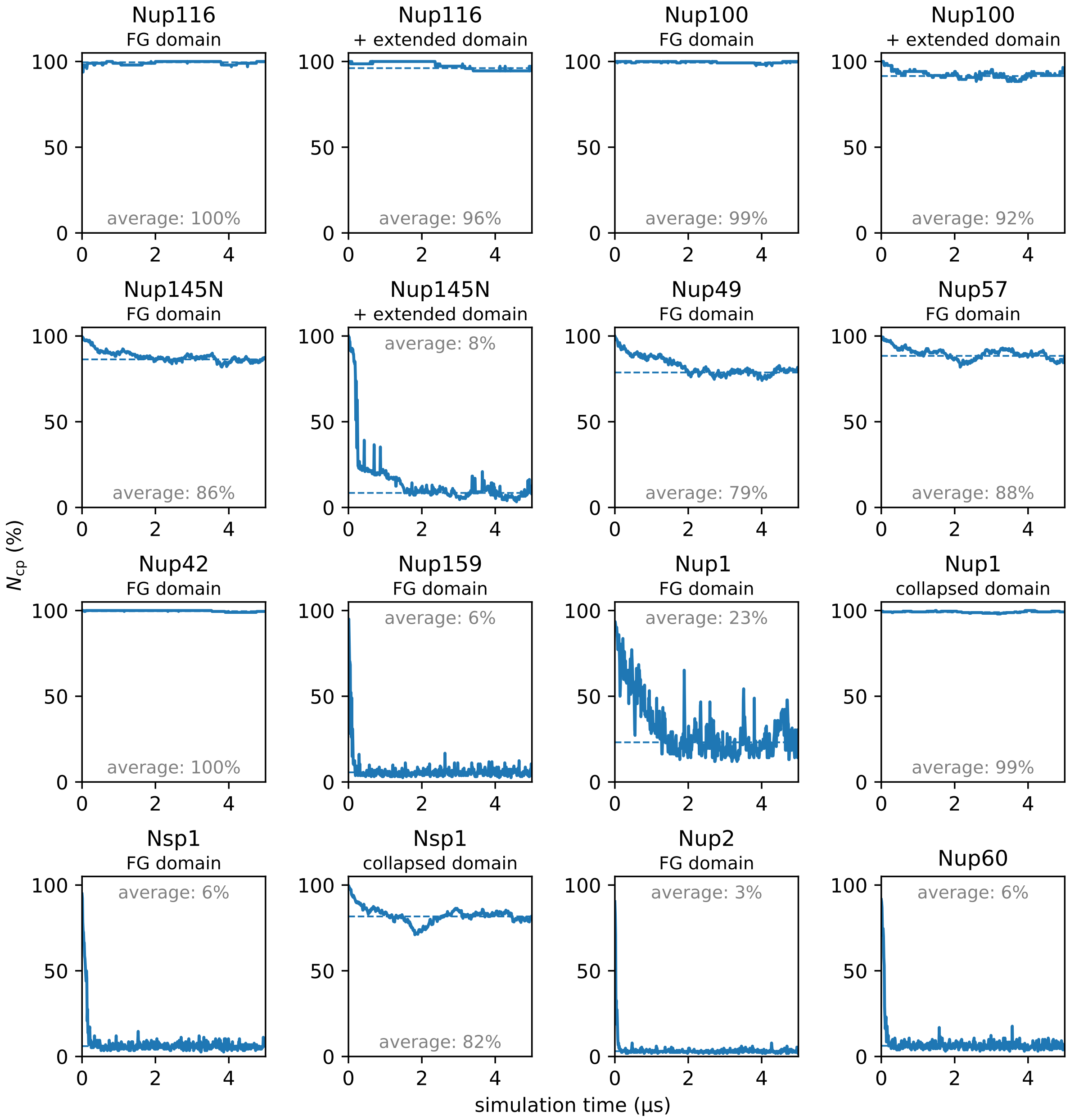

**Figure S2.** Convergence of LLPS simulations. Size of the largest cluster, $N_{cp}$ (%), as a function of simulation time for each of the LLPS simulations. Dashed lines indicated the average cluster size measured over the last 3 μs of the simulation. All simulations start from a condensate structure, therefore $N_{cp} = 100\,\%$ at $t^* = 0$. From the cluster size evolution it is clear that all simulations have reached a dynamic equilibrium within 2 μs.

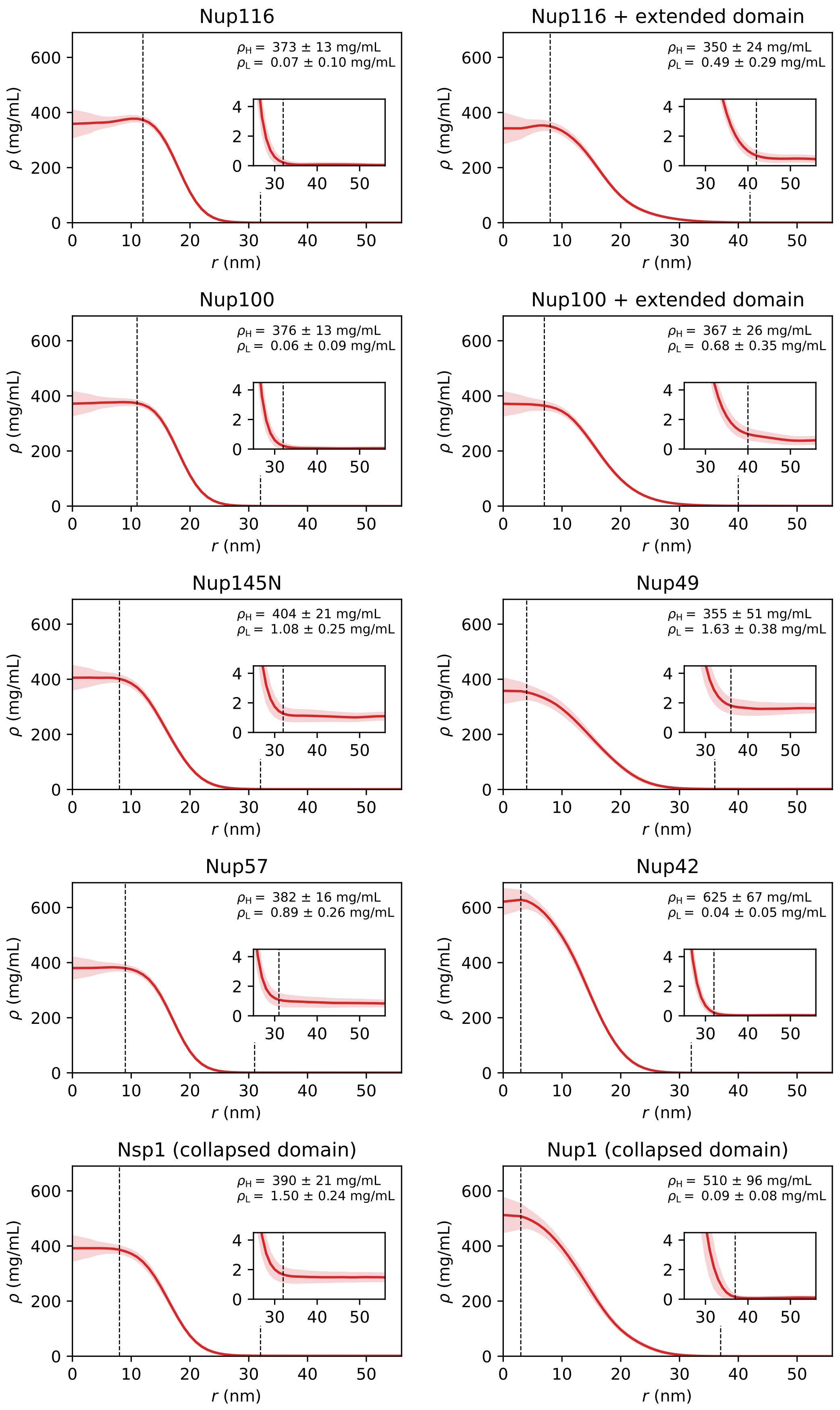

**Figure S3.** Radial density distributions for each of the FG-Nup condensates. The shading indicates half of the standard deviation as error bars. The boundaries of the high-density and low-density regions are indicated by dashed lines. The inset figures show the zoomed in density profiles for $r \geq 25\,\text{nm}$ to highlight the boundary of the low-density region.

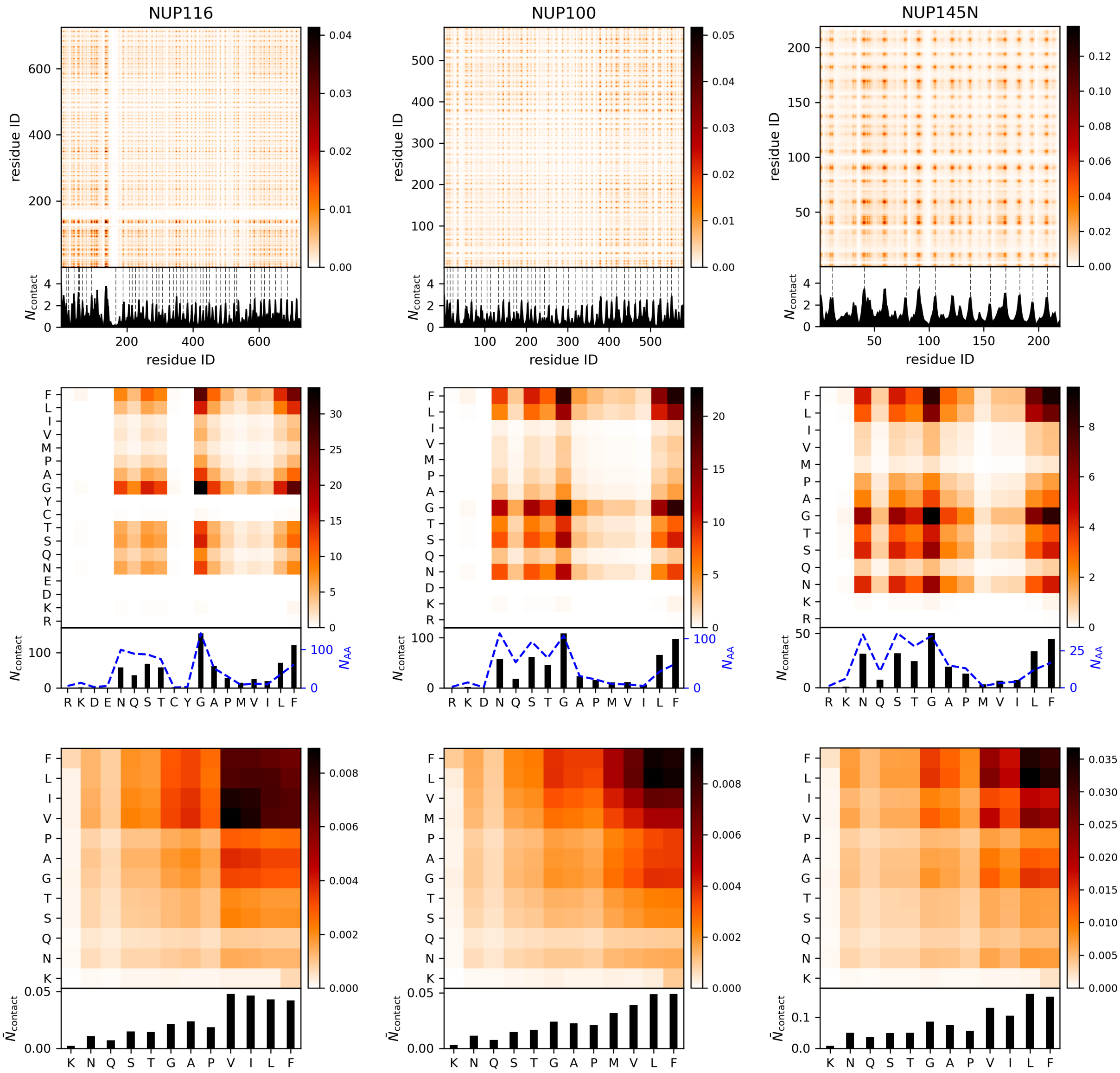

**Figure S4.** Intermolecular contact maps of the FG-Nup condensates of Nup116, Nup100 and Nup145N. **(top)** Average number of intermolecular contacts per protein replica as a function of residue number. The bottom figures show the one-dimensional summation, where the dashed lines indicate the location of the FG-motifs. **(middle)** Average number of intermolecular contacts per protein replica as a function of residue type. **(bottom)** Average number of intermolecular contacts per protein replica as a function of residue type, normalized for residue occurence in the protein sequence.

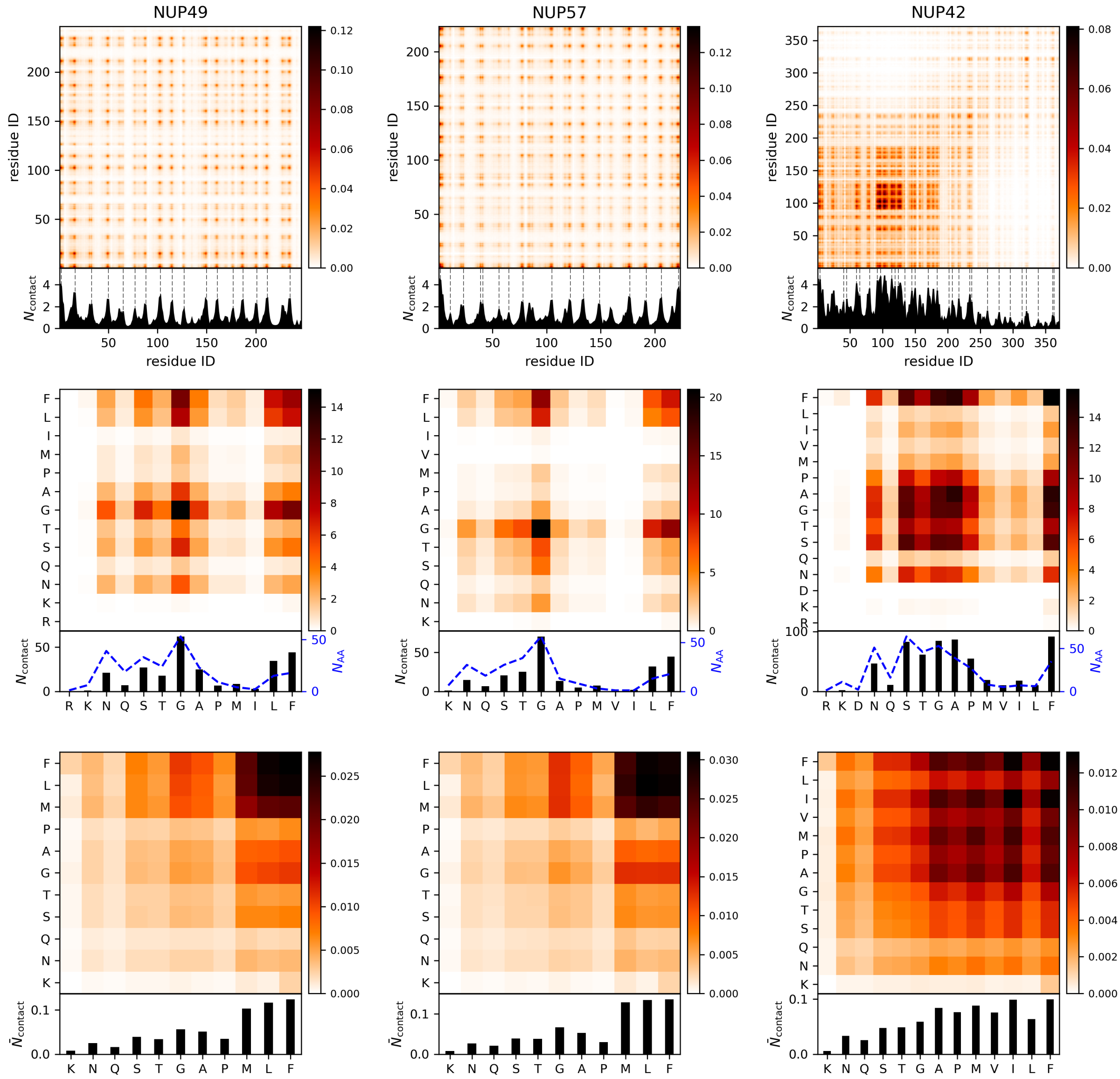

**Figure S5.** Intermolecular contact maps of the FG-Nup condensates of Nup49, Nup57 and Nup42. **(top)** Average number of intermolecular contacts per protein replica as a function of residue number. The bottom figures show the one-dimensional summation, where the dashed lines indicate the location of the FG-motifs. **(middle)** Average number of intermolecular contacts per protein replica as a function of residue type. **(bottom)** Average number of intermolecular contacts per protein replica as a function of residue type, normalized for residue occurence in the protein sequence.

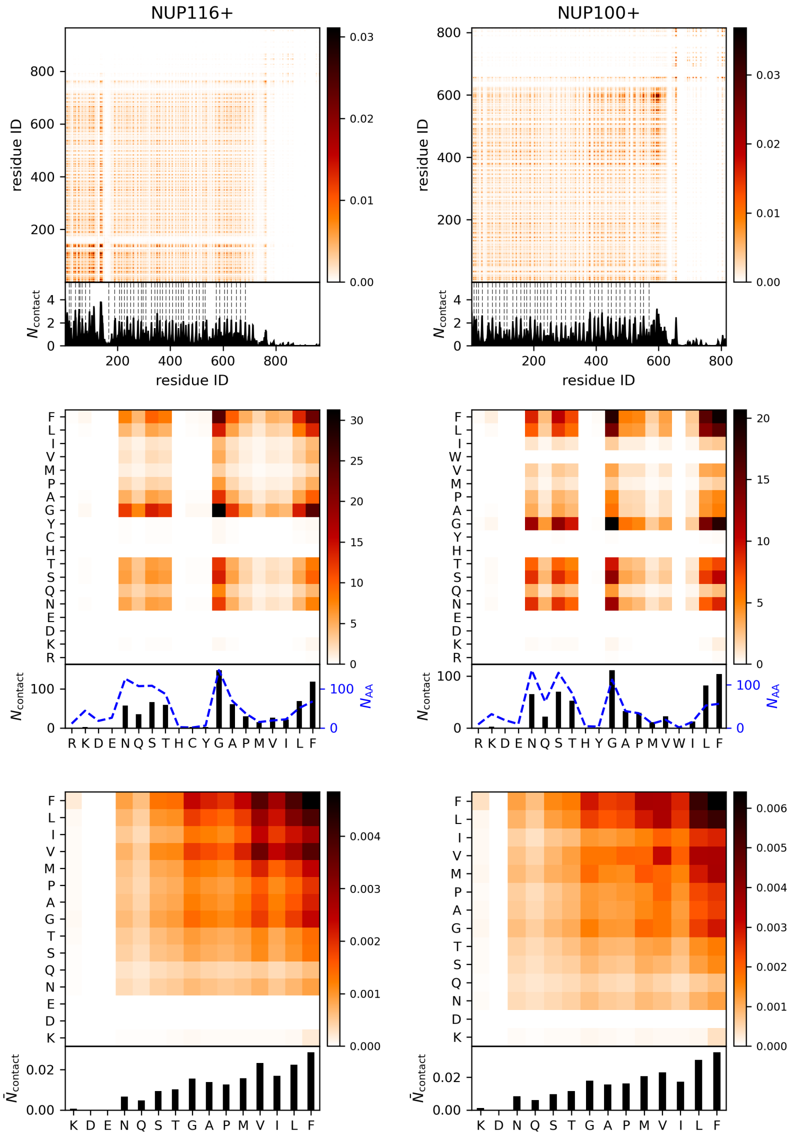

**Figure S6.** Intermolecular contact maps of the FG-Nup condensates of Nup116+ and Nup100+. **(top)** Average number of intermolecular contacts per protein replica as a function of residue number. The bottom figures show the one-dimensional summation, where the dashed lines indicate the location of the FG-motifs. **(middle)** Average number of intermolecular contacts per protein replica as a function of residue type. **(bottom)** Average number of intermolecular contacts per protein replica as a function of residue type, normalized for residue occurence in the protein sequence.

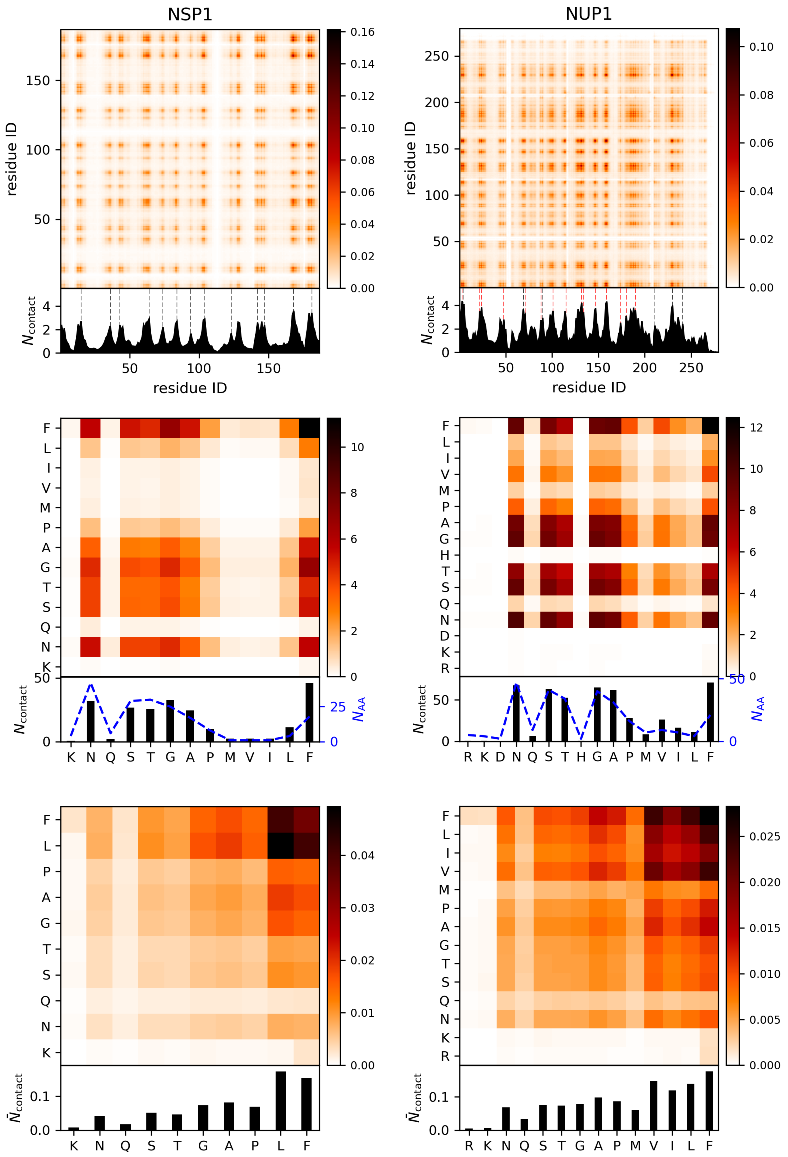

**Figure S7.** Intermolecular contact maps of the FG-Nup condensates of the Nsp1 and Nup1 (collapsed domains). **(top)** Average number of intermolecular contacts per protein replica as a function of residue number. The bottom figures show the one-dimensional summation, where the black dashed lines indicate the location of the FG-motifs. For the collapsed domain of Nup1, the location of F residues are indicated by red dashed lines. **(middle)** Average number of intermolecular contacts per protein replica as a function of residue type. **(bottom)** Average number of intermolecular contacts per protein replica as a function of residue type, normalized for residue occurence in the protein sequence.

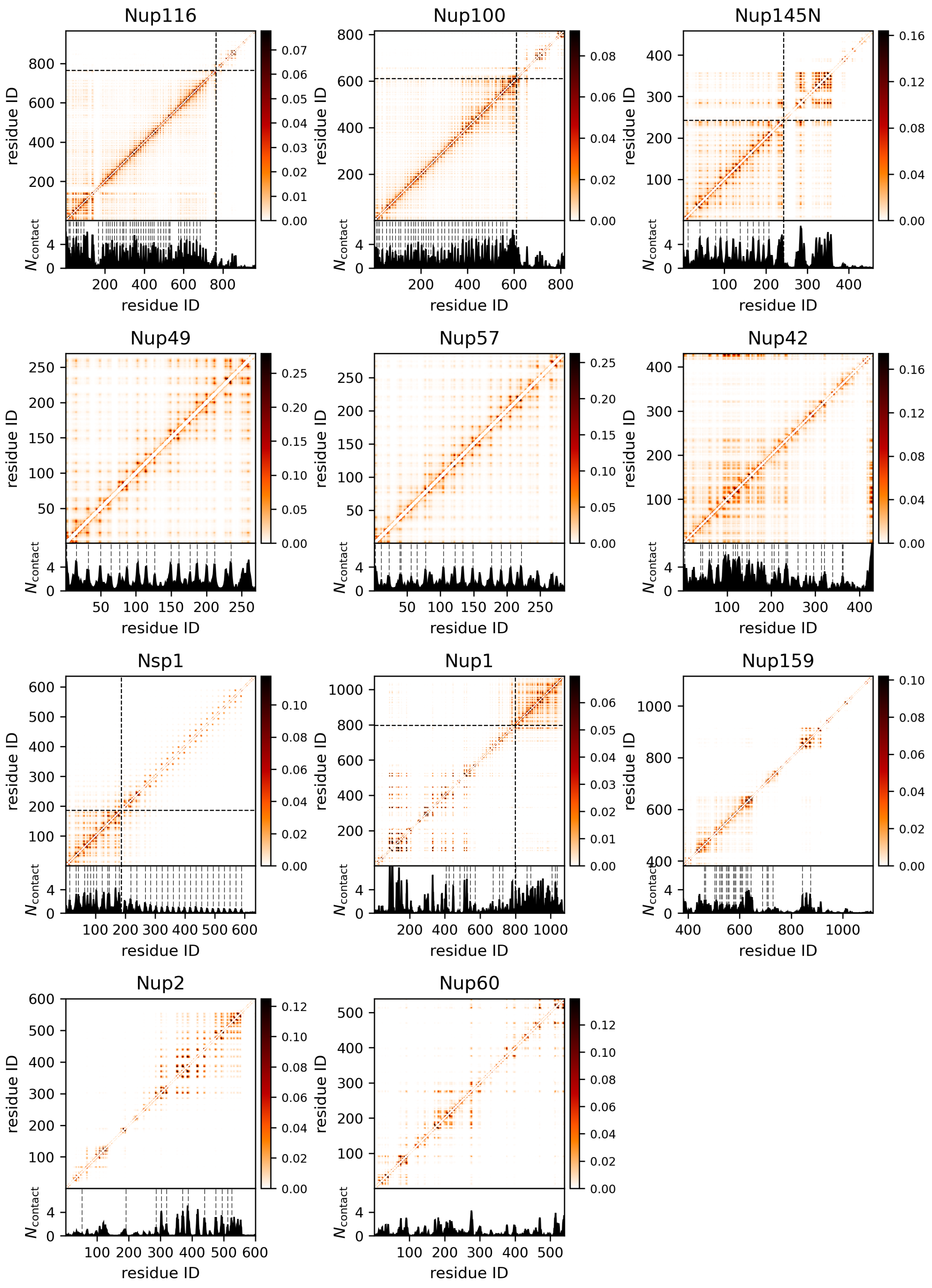

**Figure S8.** Intramolecular contact maps for each of the disordered FG-Nups. For each FG-Nup the full disordered domain (see Figure 1) is simulated. The bottom figures show the one-dimensional summation, where the dashed lines indicate the location of the FG-motifs. For the bimodal FG-Nups, i.e. Nup116, Nup100, Nup145N, Nup1 and Nsp1, a dashed line marks the boundary between the collapsed and the extended domain.

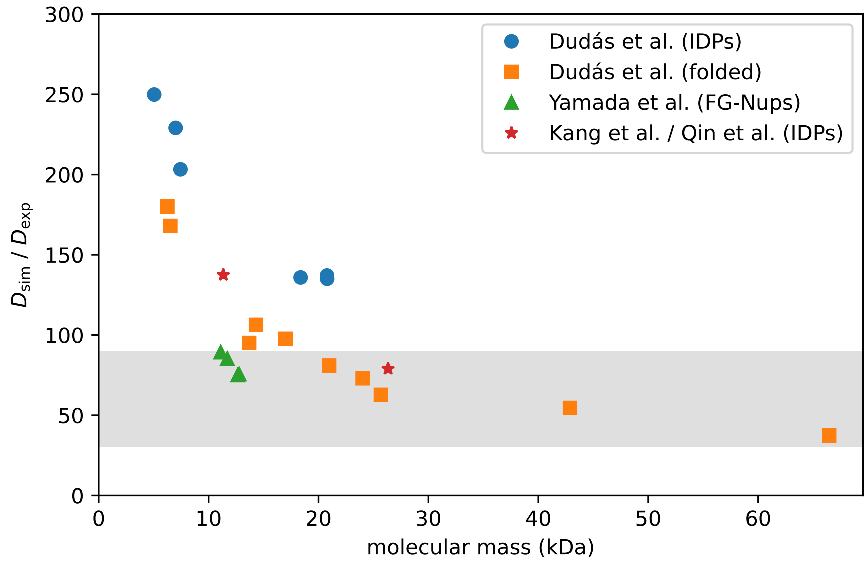

**Figure S9.** CG dynamics speed-up in 1-BPA simulations as a function of molecular mass. The speed-up factor is determined by the relative self-diffusion coefficients in our simulations and experiments (i.e. $D_{sim}/D_{exp}$). Speed-up factors are determined for a list of IDPs and folded proteins (Dudás et al. [10]), FG-Nup segments (Yamada et al. [11]), TDP-43 (Qin et al. [12]) and FUS NTD (Kang et al. [13]). We suggest that the dynamics speed-up in our simulations is in the range 30–90 times (highlighted in gray).

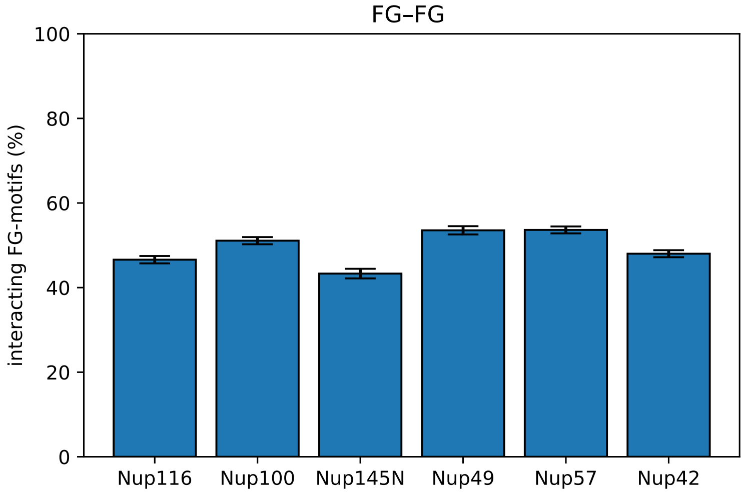

**Figure S10.** Percentage of FG-motifs involved in FG–FG interactions in FG-Nup condensates. The FG-occupancy is determined by counting the number of FG-motifs inside the FG-Nup condensate that make at least one contact with another FG-motif ($r_{\mathrm{cut}} = 0.7\,\mathrm{nm}$).

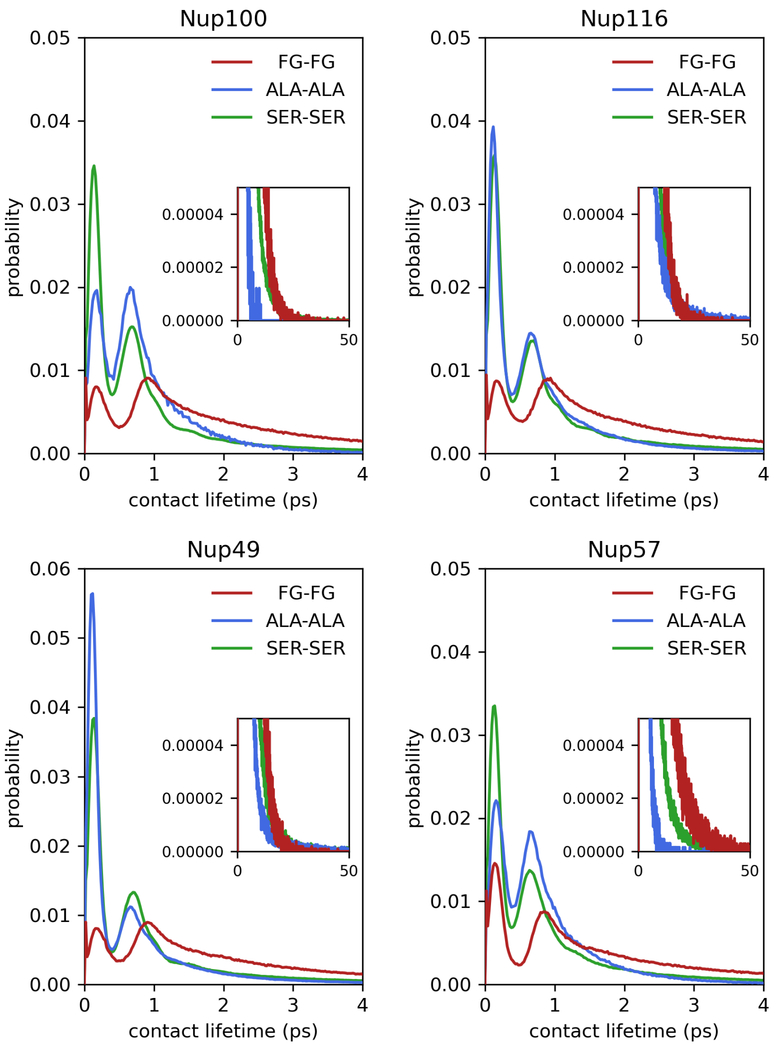

**Figure S11.** Intermolecular contact lifetimes in FG-Nups condensates ($r_c = 0.6\,\text{nm}$). Sampling is done over a trajectory of 10 ns.

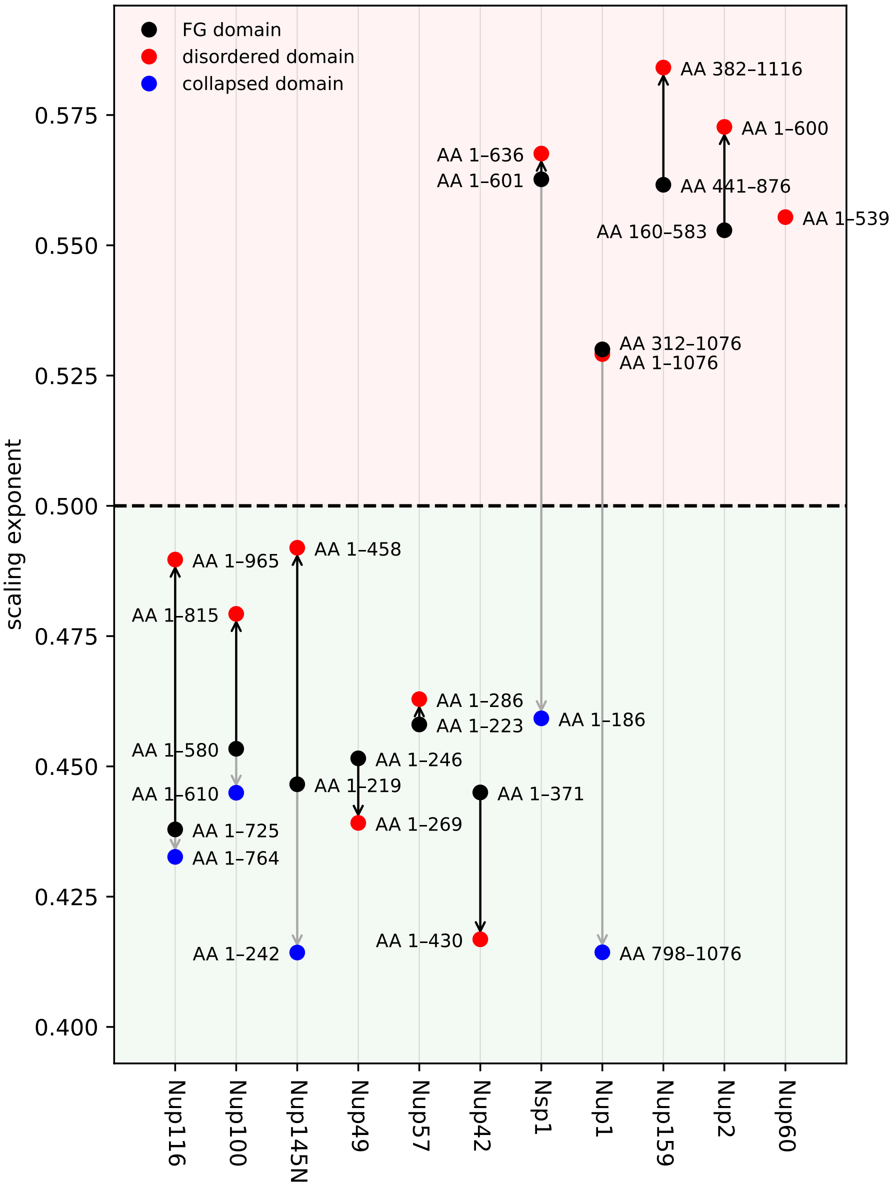

**Figure S12.** Scaling exponents for different domains of each FG-Nup: FG domains used in the LLPS simulations (black), full disordered domains based on the data of Kim et al. [14] (red), and collapsed domains as described by Yamada et al. [11] (blue). In general, LLPS is observed for scaling exponents $< 0.5$.

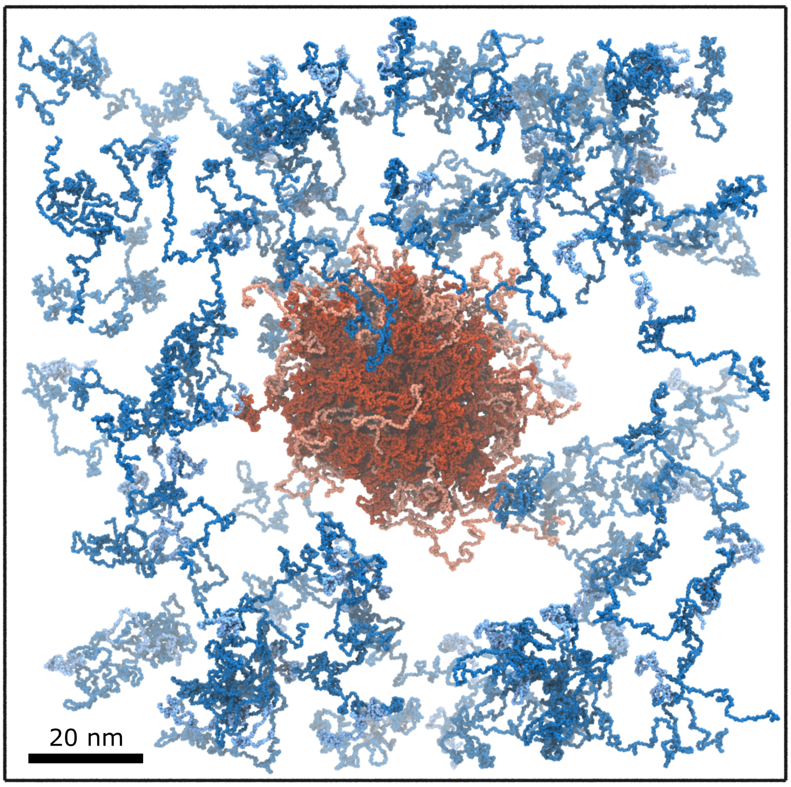

**Figure S13.** Initial configuration of the NPC condensate simulation. FG-Nups with a scaling exponent $\nu < 0.5$ (red, with the extended domains of Nup116, Nup100 and Nup145N in pink) are in the condensate structure are colored in red. FG-Nups with a scaling exponent $\nu > 0.5$ (blue, with the collapsed domains of Nsp1 and Nup1 colored in light blue) are randomly distributed around the condensate.

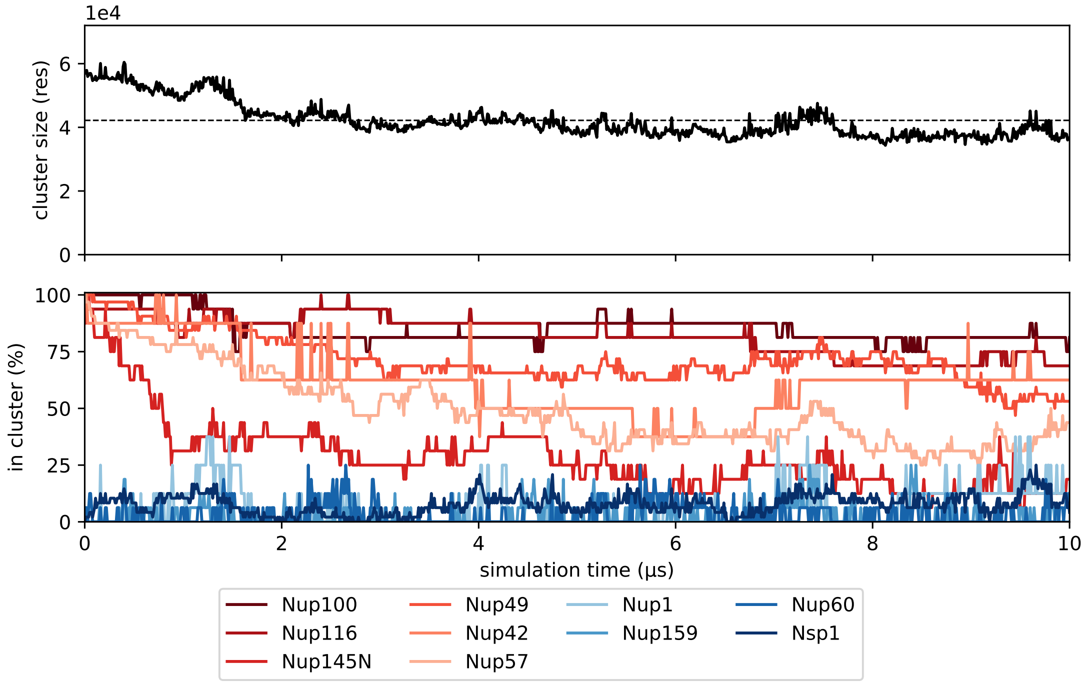

**Figure S14.** Convergence of the NPC condensate simulation. (top) Size of the NPC condensate (in number of residues) as a function of simulation time. The dashed line indicates the average cluster size over the last 5 µs of the simulation. (bottom) Composition of the NPC condensate as a function of simulation time. For each FG-Nup type, the percentage of monomers that is present in the NPC condensate is shown over time. From the cluster size evolution it is clear that the simulation has reached a dynamic equilibrium within 5 µs. The equilibrated NPC condensate contains Nup100 (84 %), Nup116 (75 %), Nup49 (66 %), Nup42 (54 %), Nup57 (37 %), Nup145N (18 %) and a small fraction of Nsp1 (9 %).

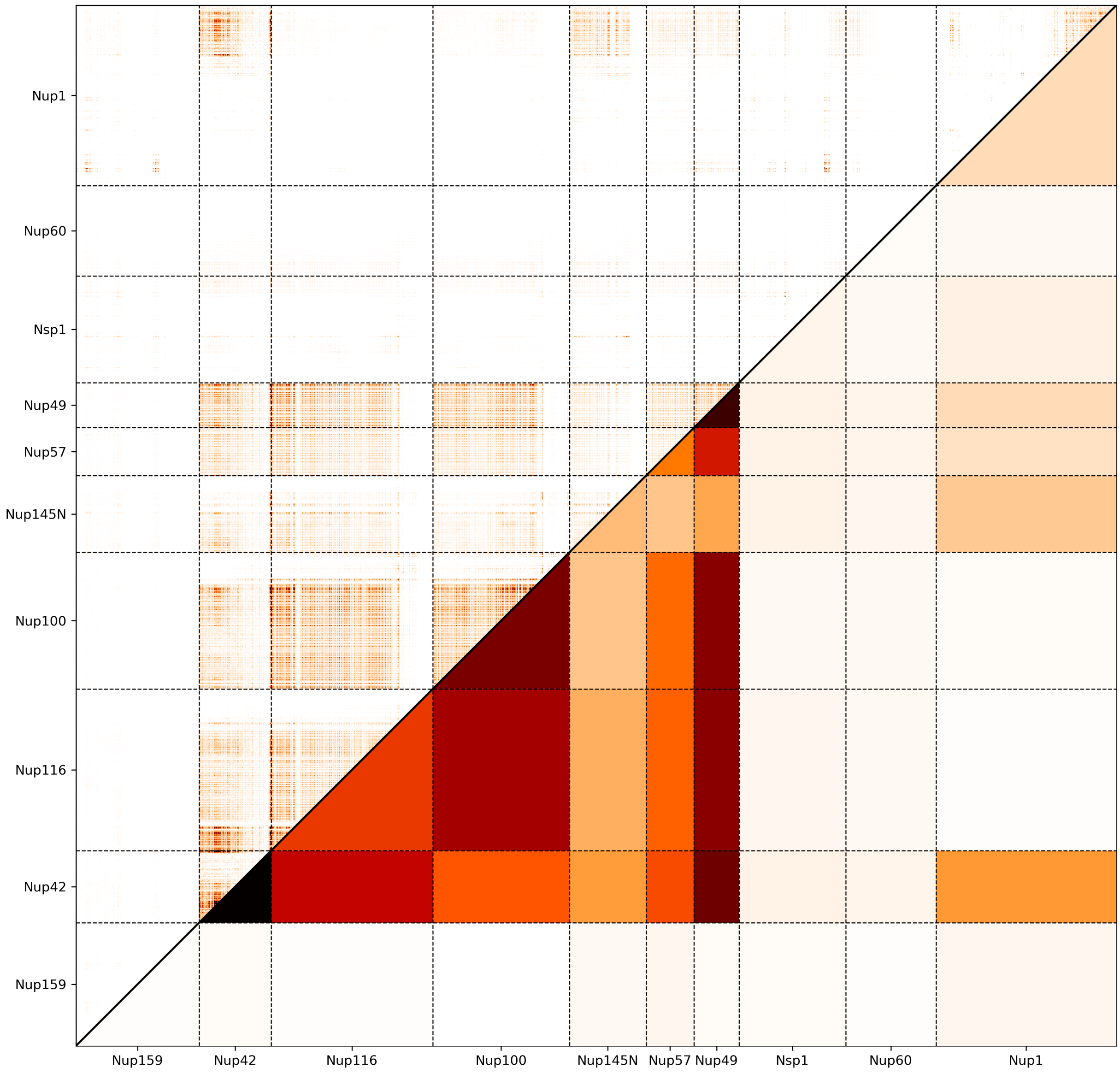

**Figure S15.** Intermolecular contact analysis of the NPC condensate simulation. The upper triangle shows the average number of intermolecular contacts per protein replica as a function of residue number for each of the FG-Nup combinations. The lower triangle shows the average protein contacts per protein replica. Contact numbers are normalized for FG-Nup abundance.

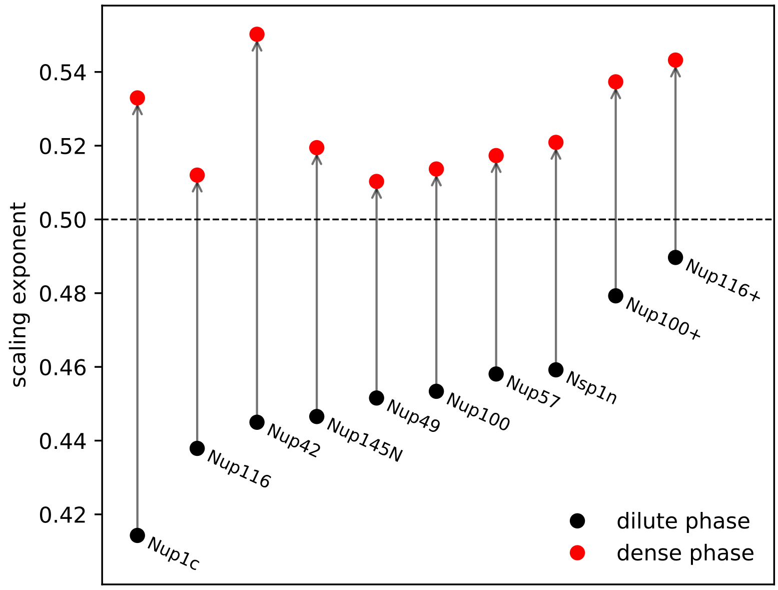

**Figure S16.** Comparison of FG-Nup scaling exponents in the dilute phase and the dense phase. Scaling exponents of the dilute phase are obtained from single-chain simulations (see Figure S12). Scaling exponents of the dense phase are obtained by determining the radius of gyration of the FG-Nups inside the FG-Nup condensate, averaged over the last 2.5 μs of each simulation.

### 5 Supplementary Tables

| amino acid | mass (Da) | $\varepsilon_i$ | charge ($e$) | amino acid | mass (Da) | $\varepsilon_i$ | charge ($e$) |
|---|---|---|---|---|---|---|---|
| A | 71.07 | 0.7 | 0 | L | 113.16 | 1 | 0 |
| R | 156.19 | 0.005* | +1 | K | 128.17 | 0.005* | +1 |
| N | 114.10 | 0.41† | 0 | M | 131.20 | 0.78 | 0 |
| D | 115.09 | 0.005* | −1 | F | 147.18 | 1 | 0 |
| C | 103.14 | 0.68 | 0 | P | 97.12 | 0.65 | 0 |
| Q | 128.13 | 0.33† | 0 | S | 87.08 | 0.45 | 0 |
| E | 129.11 | 0.005* | −1 | T | 101.11 | 0.51 | 0 |
| G | 57.05 | 0.48† | 0 | W | 186.22 | 0.96 | 0 |
| H | 137.14 | 0.53 | 0 | Y | 163.18 | 0.82 | 0 |
| I | 113.16 | 0.98 | 0 | V | 99.13 | 0.94 | 0 |

**Table S1.** Relative hydrophobic strength $\varepsilon_i$ and charge for each amino acid [2]. *The hydrophobicity values of the charged residues have been slightly increased in line with recent work [15]. †The hydrophobicity values of G, Q and N have been updated such that the 1-BPA force field better reproduces experimental data of Yamada et al. [11].

| Cation–$\pi$ pair | Energy (kJ/mol) |
|---|---|
| R – F | 4.30 |
| R – Y | 5.00 |
| R – W | 6.70 |
| K – F | 1.79 |
| K – Y | 3.13 |
| K – W | 4.26 |

**Table S2.** Cation–$\pi$ interaction energies, $\varepsilon_{\mathrm{cp},ij}$, for all six cation–$\pi$ pairs [3].

| FG-Nup segment | $R_{h,exp}$ | 1-BPA-cp (old) $R_{h,sim}$ | error (%) | 1-BPA-v2 (new) $R_{h,sim}$ | error (%) |
|---|---|---|---|---|---|
| Nsp1n_lc | 27.1 | 32.8 | 20.9 | 29.2 | 7.7 |
| Nup116m_lc | 46.5 | 45.9 | 1.4 | 46.2 | 0.5 |
| Nup100n_lc | 48.7 | 49.1 | 0.9 | 47.3 | 2.9 |
| Nup49_lc | 26.9 | 33.1 | 23.1 | 30.5 | 13.4 |
| Nup42_lc | 28.4 | 29.4 | 3.6 | 27.7 | 2.3 |
| Nup57_lc | 31.9 | 35.0 | 9.8 | 34.2 | 7.2 |
| Nup145N_lc | 28.2 | 32.0 | 13.4 | 30.1 | 6.7 |
| Nup1c_lc | 32.4 | 36.0 | 11.1 | 32.4 | 0.1 |
| Nup159_hc | 55.4 | 59.1 | 6.7 | 57.4 | 3.6 |
| Nup60_hc | 31.3 | 34.0 | 8.6 | 34.1 | 8.9 |
| Nup1m_hc | 67.9 | 68.1 | 0.3 | 67.3 | 0.8 |
| Nup2_hc | 59.8 | 54.0 | 9.8 | 52.1 | 12.9 |
| Nsp1m_hc | 65.3 | 63.8 | 2.3 | 62.7 | 4.0 |
| Nup145Ns | 29.8 | 36.1 | 21.1 | 36.1 | 21.1 |
| Nup100s | 36.6 | 40.6 | 10.9 | 40.2 | 9.8 |
| Nup116s | 39.1 | 42.6 | 8.9 | 42.6 | 9.0 |
| **average** | | | **9.5** | | **6.9** |

**Table S3.** Stokes radii of all Yamada FG-Nup segments for the original 1-BPA model (with cation-$\pi$) and the updated hydrophobicities of glycine, glutamine and asparagine. All single-chain simulations are performed at $T = 303\,\mathrm{K}$ and salt concentration 150 mM ($\kappa = 1.27$). Errors are calculated as $|R_{h,sim} - R_{h,exp}|/R_{h,exp}$.

| FG-Nup | segment | LLPS | experiment | comment |
|---|---|---|---|---|
| Nup116 | AA 165–716 | yes | Patel et al. [16] | low-affinity binding interactions |
| Nup116 | AA 165–716 | yes | Patel et al. [16] | in-vivo |
| Nup116 | AA 1–736 | yes | Schmidt & Görlich [17] | hydrogels |
| Nup116 | AA 1–725 | yes | Kuiper et al. [18] | aggregates |
| Nup116 | AA 1–725 | yes | this work | |
| Nup116 | AA 1–965 | yes | this work | |
| Nup100 | AA 1–640 | yes | Patel et al. [16] | low-affinity binding interactions |
| Nup100 | AA 1–640 | yes | Patel et al. [16] | in-vivo |
| Nup100 | AA 1–580 | yes | Schmidt & Görlich [17] | hydrogels |
| Nup100 | AA 1–580 | yes | Kuiper et al. [18] | aggregates |
| Nup100 | AA 1–580 | yes | this work | |
| Nup100 | AA 1–815 | yes | this work | |
| Nup145N | AA 1–262 | yes | Patel et al. [16] | in-vivo |
| Nup145N | AA 1–219 | yes | Kuiper et al. [18] | aggregates |
| Nup145N | AA 1–219 | yes | this work | |
| Nup145N | AA 1–458 | no | this work | |
| Nup49 | AA 1–236 | yes | Patel et al. [16] | in-vivo |
| Nup49 | AA 1–260 | yes | Milles et al. [19] | hydrogels |
| Nup49 | AA 1–249 | yes | Celetti et al. [20] | |
| Nup49 | AA 1–246 | yes | this work | |
| Nup57 | AA 1–255 | yes | Patel et al. [16] | low-affinity binding interactions |
| Nup57 | AA 1–255 | yes | Patel et al. [16] | in-vivo |
| Nup57 | AA 1–223 | yes | this work | |
| Nup42 | AA 29–129 | yes | Patel et al. [16] | low-affinity binding interactions |
| Nup42 | AA 1–364 | (?) | Patel et al. [16] | in-vivo |
| Nup42 | AA 1–371 | yes | this work | |
| Nup159 | AA 433–879 | no | Patel et al. [16] | low-affinity binding interactions |
| Nup159 | AA 433–879 | no | Patel et al. [16] | in-vivo |
| Nup159 | AA 441–876 | no | this work | |
| Nup1 | AA 332–1076 | no | Patel et al. [16] | low-affinity binding interactions |
| Nup1 | AA 312–1076 | no | this work | |
| Nup1 | AA 798–1076 | yes | this work | |
| Nsp1 | AA 1–601 | yes | Frey et al. [21] | hydrogels |
| Nsp1 | AA 1–603 | no | Patel et al. [16] | low-affinity binding interactions |
| Nsp1 | AA 1–591 | no | Patel et al. [16] | in-vivo |
| Nsp1 | AA 1–603 | no | Yamada et al. [11] | low-affinity binding interactions |
| Nsp1 | AA 1–601 | no | this work | |
| Nsp1 | AA 1–172 | yes | Yamada et al. [11] | low-affinity binding interactions |
| Nsp1 | AA 1–186 | yes | this work | |
| Nsp1 | AA 173–603 | no | Yamada et al. [11] | low-affinity binding interactions |
| Nsp1 | AA 274–601 | no | Schmidt & Görlich [17] | hydrogels |
| Nup2 | AA 181–537 | (?) | Patel et al. [16] | low-affinity binding interactions |
| Nup2 | AA 160–583 | no | this work | |
| Nup60 | AA 387–521 | no | Patel et al. [16] | in-vivo |
| Nup60 | AA 1–539 | no | this work | |

**Table S4.** Overview of LLPS of yeast FG-Nups in experiments.

| FG-Nup | AA domain | length | predominant FG type | number of FG-motifs | charged AAs (RKDE) [%] | hydrophobic AAs (ILFYWV) [%] | C/H ratio | scaling exp. | LLPS |
|---|---|---|---|---|---|---|---|---|---|
| Nup116 | 1–725 | 725 | GLFG | 47 | 3 | 16 | 0.21 | 0.44 | yes |
| Nup116+ | 1–965 | 965 | GLFG | 47 | 10 | 17 | 0.60 | 0.49 | yes |
| Nup100 | 1–580 | 580 | GLFG | 44 | 2 | 16 | 0.15 | 0.45 | yes |
| Nup100+ | 1–815 | 815 | GLFG | 44 | 8 | 18 | 0.46 | 0.48 | yes |
| Nup145N | 1–219 | 219 | GLFG | 11 | 3 | 16 | 0.19 | 0.45 | yes |
| Nup145N+ | 1–458 | 458 | GLFG | 11 | 13 | 19 | 0.72 | 0.49 | no |
| Nup49 | 1–246 | 246 | GLFG | 17 | 3 | 14 | 0.20 | 0.45 | yes |
| Nup57 | 1–223 | 223 | GLFG | 16 | 3 | 15 | 0.18 | 0.46 | yes |
| Nup42 | 1–371 | 371 | SAFGxPSFG | 29 | 4 | 14 | 0.26 | 0.45 | yes |
| Nup159 | 441–876 | 436 | SAFGxPSFG | 25 | 18 | 17 | 1.01 | 0.56 | no |
| Nup1 | 312–1076 | 765 | FxFG | 17 | 18 | 16 | 1.16 | 0.53 | no* |
| Nup1c | 798–1076 | 279 | FG | 6 | 4 | 15 | 0.27 | 0.41 | yes |
| Nsp1 | 1–601 | 601 | FxFG | 33 | 22 | 12 | 1.90 | 0.56 | no |
| Nsp1n | 1–186 | 186 | FG | 13 | 2 | 13 | 0.17 | 0.46 | yes |
| Nup2 | 160–583 | 424 | FxFG | 11 | 27 | 15 | 1.74 | 0.55 | no |
| Nup60 | 1–539 | 539 | FxFx | 0 | 25 | 22 | 1.14 | 0.56 | no |

**Table S5.** Properties of the FG-Nup domains of yeast.
